# Restriction-weighted q-space trajectory imaging (ResQ): Toward mapping diffusion-time effects with tensor-valued diffusion encoding in human prostate cancer xenografts

**DOI:** 10.64898/2025.12.08.692924

**Authors:** Filip Szczepankiewicz, Malwina Molendowska, Samo Lasič, Marcella E Safi, Michael Gottschalk, Evangelia Sereti, Anders Bjartell, Linda Knutsson, Oskar Vilhelmsson Timmermand, Crister Ceberg, Joanna Strand

**Affiliations:** Department of Medical Radiation Physics, Lund University, Lund, Sweden; Department of Diagnostic Radiology, Lund University, Lund, Sweden; Department of Clinical Sciences Lund, Lund University, Lund, Sweden; Lund University Bioimaging Centre, Lund University, Lund, Sweden; Department of Translational Medicine, Lund University, Malmö, Sweden; Department of Urology, Skåne University Hospital, Malmö, Sweden; Department of Clinical Sciences Lund, Oncology, Lund University, Lund, Sweden; Department of Hematology, Oncology, Radiation Physics, Skåne University Hospital, Lund University, Lund, Sweden

## Abstract

**Purpose:** Tensor-valued diffusion encoding employs gradient waveforms that enable unique sensitivity to microstructural features of tissue, but the interpretation of signal and parameters may be confounded by diffusion-time dependence. We introduce a framework for restriction-weighted q-space trajectory imaging (ResQ) that incorporates diffusion-time effects via the restriction-weighting tensor, and we evaluate it in a longitudinal study of prostate cancer xenografts treated by external radiotherapy.

**Methods:** We proposed a novel gradient waveform design for tensor-valued encoding with controlled restriction weighting and applied a set of four waveforms at a 9.4 T preclinical MRI system. Mice were inoculated with human prostate cancer cells (LNCaP) and assigned to groups that were untreated controls or treated by external beam irradiation. ResQ produced parameters that describes the diffusion process in terms of the mean diffusivity (*D*), isotropic diffusional variance (*V*_Di_), and microscopic diffusion anisotropy (*V*_Da_) as well as their diffusion-time dependence (Δ*D*, Δ*V*_Di_, Δ*V*_Da_). Analyses were performed to characterize parameters longitudinally and across groups. To highlight the consequences of ignoring restriction effects, we compared ResQ to analogous parameters estimated by q-space trajectory imaging (QTI).

**Results:** ResQ revealed clear diffusion-time dependence across all tumors, with significant longitudinal differences between treated and untreated groups, most prominent in *D*, Δ*D*, and *V*_Di_. The ResQ signal representation captured the signal dynamics, whereas QTI did not. Neglecting diffusion-time dependence in QTI led to substantial parameter bias, most notably a pronounced overestimation of microscopic diffusion anisotropy.

**Conclusion:** Diffusion-time effects are non-negligible in prostate cancer and must be considered when using tensor-valued diffusion encoding. The ResQ framework enables controlled restriction weighting and improved interpretability of diffusion MRI parameters compared to approaches that ignore the effects of restriction. This provides a more principled approach for tensor-valued diffusion encoding and may enable novel imaging biomarkers that disentangle diffusivity, isotropic diffusional variance, microscopic anisotropy, and their diffusion-time dependence.

## Introduction

Diffusion MRI (dMRI) is uniquely sensitive to features of tissue microstructure and can probe them noninvasively without exogenous contrast agents (Fokkinga et al., 2023). It relies on the principle that subtle alterations in the tissue microstructure caused by, for example, cancer growth and progression, perturb the diffusive motion of water molecules in a manner that is detectable and quantifiable by dMRI. Indeed, a prominent clinical application of dMRI is in examining patients who are suspected of having prostate cancer (Weinreb et al., 2016, Tempany et al., 2015). However, current diagnostic routine relies on relatively simplistic dMRI methods that fail to detect a relevant fraction of lesions (Chatterjee and Dwivedi, 2024, Nam et al., 2022, Padhani et al., 2019) and struggle to distinguish different tumor types that may lead to inaccurate staging and tumor grading (Agrotis et al., 2025, Chatterjee et al., 2023).

In recent years, several methodologies have been proposed for prostate cancer imaging which aim to improve earlier detection, accurate staging, guidance for biopsies, and planning/monitoring treatment (Zhu et al., 2024, Molendowska et al., 2024, Chatterjee and Dwivedi, 2024, Singh et al., 2022, Zhang et al., 2020, Johnston et al., 2019, Hectors et al., 2018, Stoyanova et al., 2018). Common among them is the use of specialized measurements and analysis methods that go beyond measuring the diffusion-weighted signal and apparent diffusion coefficient. The driving principle is that ever more subtle features of microstructure can be detected or quantified when an appropriate combination of measurement dimensions, including relaxation weighting, is probed jointly in a ‘multidimensional’ or ‘hybrid’ fashion (Benjamini and Basser, 2020, Lundell et al., 2019, de Almeida Martins and Topgaard, 2018, Chatterjee et al., 2018, Westin et al., 2016, Wang et al., 2014, Novikov et al., 2018, Lampinen et al., 2023).

An especially promising family of multidimensional methods uses so-called tensor-valued diffusion encoding, wherein non-conventional gradient waveforms are used to probe the diffusion process along more than one direction per shot. This enables investigation of features that are not accessible by conventional diffusion encoding, such as microscopic diffusion anisotropy and isotropic diffusion heterogeneity (Lasič et al., 2014, Finsterbusch, 2011, Jespersen et al., 2013, Shemesh et al., 2010, Cory et al., 1990). For example, ‘diffusional variance decomposition’ (Lasič et al., 2014, Szczepankiewicz et al., 2016) and ‘q-space trajectory imaging’ (Westin et al., 2016), have been used to distinguish tumor types and grades based on their diffusion parameters (Cho et al., 2024, Szczepankiewicz et al., 2016, Brabec et al., 2022), as well as multiple preliminary studies in prostate cancer (Langbein et al., 2021, Nilsson et al., 2021, Molendowska et al., 2024, Szczepankiewicz et al., 2023).

Rather than using the conventional Stejskal-Tanner encoding (Stejskal and Tanner, 1965), i.e., a pair of trapezoidal pulsed field gradients, the encoding is performed with continuous modulation of the gradient waveform. The latter is called ‘free waveform encoding’ as it enables a highly flexible framework for experimental design with superior encoding efficiency, i.e., shorter encoding times for a given b-value compared to a series of trapezoids (Szczepankiewicz et al., 2021). However, the temporal modulation of the gradient also affects the diffusion time, which may vary within and between the employed gradient waveforms (de Swiet and Mitra, 1996, Szczepankiewicz et al., 2021, Jespersen et al., 2018, Lasič et al., 2025). In the case of Gaussian diffusion, which is independent of diffusion time, it has no effect on the result and can be safely ignored. Conversely, if time-dependence is present, a non-trivial interaction between the gradient waveform and the underlying microstructure geometry can confound both quantification and interpretation of the estimated diffusion parameters (Jespersen et al., 2018, Lundell et al., 2019).

The microstructure in healthy prostate and prostate cancer is such that diffusion-time effects are likely to arise in most practical diffusion-weighted measurements; it is a highly heterogeneous tissue that comprises restrictions with a wide range of sizes (Yadav et al., 2018) and anisotropic stromal tissue (Bourne et al., 2016). Indeed, time-dependent diffusion has been reported in both healthy prostate (Ma et al., 2025) and prostate cancer (Lemberskiy et al., 2017, Wu et al., 2022), and it serves as a pivotal mechanism for generating biomarkers in several dMRI methods (Panagiotaki and Alexander, 2014, Jiang et al., 2017). As stated in preliminary studies of prostate (Langbein et al., 2021, Nilsson et al., 2021), diffusion-time effects are likely to confound the results, which could explain why there was no clear relation between the diffusion parameters and the qualitative analysis of histology. Rather than avoiding tensor-valued diffusion encoding in these challenging scenarios, we strive to incorporate diffusion-time effects in the signal representation so that the restriction weighting can be controlled and become a valuable imaging biomarker rather than a confounder.

In this study, we investigated the effects of diffusion time and restrictions in a longitudinal study in mice that carry human prostate cancer xenografts and undergo treatment by external beam irradiation. We propose a novel design for tensor-valued diffusion encoding with controlled restriction weighting, and we use it to establish that restricted diffusion has a marked impact on the diffusion-weighted signal and estimated parameters. To facilitate the analysis, we propose a methodological framework for restriction-weighted q-space trajectory imaging (ResQ); a phenomenological representation of the diffusion process that leverages the effects of restriction as potential microstructure biomarkers.

## Theory

In this section, we present a theoretical framework that accounts for heterogeneity, micro-anisotropy, and restrictions in a substrate composed of multiple compartments or domains, with the overall aim of describing the effects of restrictions in tissue. Since we perform non-linear diffusion encoding with arbitrarily modulated gradient waveforms, the method is closely related to q-space trajectory imaging (Westin et al., 2016) wherein effects of restrictions are not accounted for. Since arbitrary gradient modulation does not afford a straightforward definition of the diffusion time, we will employ a description of the restriction weighting that captures diffusion-time dependence via the power spectrum of the dephasing vector, as detailed below.

### q-space trajectory imaging (QTI)

Q-space trajectory imaging (Westin et al., 2016) is an extension to methods like diffusion tensor and diffusional kurtosis imaging (Basser et al., 1994, Jensen et al., 2005), that enables the estimation of microscopic diffusion anisotropy and isotropic diffusion heterogeneity by varying the ‘shape’ of the diffusion encoding. Thereby it can resolve features that are inaccessible by conventional dMRI (Lasič et al., 2014, Szczepankiewicz et al., 2016).

The signal for Gaussian diffusion in a single homogeneous environment can be described by the diffusion tensor (**D**_µ_), according to (Basser et al., 1994)

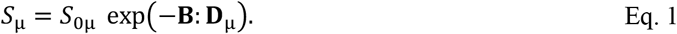

Here, **B** is the diffusion encoding b-tensor, defined as (Westin et al., 2016)

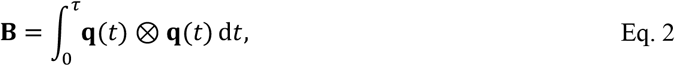

and the dephasing vector is the cumulative integral over the time-varying effective gradient waveform (**g**(*t*)), according to

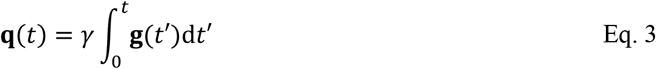

where γ is the gyromagnetic ratio, τ is the echo time, and ⊗ denotes the outer product.

In a mixture of Gaussian contributions, the powder-averaged signal can be approximated by a cumulant expansion (Kiselev, 2011, Grebenkov, 2007)

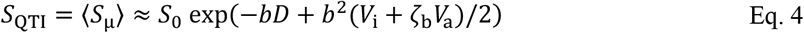

where brackets ⟨⋅⟩ indicate averaging over all domains in a voxel, *D* is the average diffusivity, and the higher order terms describe the isotropic and anisotropic diffusional variance (*V*_i_ and *V*_a_). Since we employ a ‘powder average’ representation of the signal, we may describe the encoding by the ‘size’ and ‘shape’ of the b-tensor (Topgaard, 2016). Thus, the b-value is

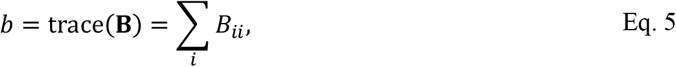

and the shape, or sensitivity to anisotropy, is

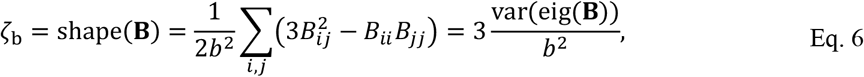

where *B*_*ij*_ denotes the elements of the b-tensor, var(⋅) is the population variance operator^1^, and eig(⋅) is the eigenvalue operator. Note that the shape parameter takes on values between 0 and 1 for spherical and linear b-tensor encoding, and carries the same information as the shape parameter introduced by Eriksson et al. (2015), where it was denoted 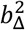. We use the current formalism because it does not require that eigenvalues are calculated, and it generalizes for non-axisymmetric encoding tensors.

### Restriction-weighted q-space trajectory imaging (ResQ)

To capture the effects of both Gaussian and restricted diffusion in a voxel that contains multiple microdomains, we generalize the QTI framework to account for variable restriction weighting using a series of approximations. We begin by describing the diffusion process in each micro-domain by its velocity autocorrelation function, or equivalently its Fourier transform, the diffusion spectrum *D*(ω) (Stepisnik, 1993, Stepisnik, 1999). Using the Gaussian phase approximation (Stepisnik, 1999), the signal in each micro-domain (*S*_µ_) is given by the spectral overlap between the encoding spectrum of an arbitrarily modulated gradient waveform and the diffusion spectrum

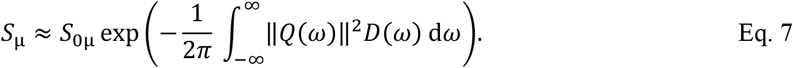

Notably, Eq. 7 omits higher order effects, such as intra-compartment kurtosis, however, the mono-exponential form does not imply free Gaussian diffusion or a flat diffusion spectrum since restriction effects enter through the frequency dependence of *D*(ω). Thus, the higher-order terms, introduced in Eq. *12* and beyond, arise from the heterogeneous distribution of domain-level apparent tensors, not from intra-domain non-Gaussian phase dispersion.

For restricted diffusion in the low-frequency regime considered here, we approximate the diffusion spectrum by a quadratic expansion (Stepisnik, 1999, Nilsson et al., 2017)

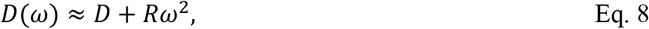

where *D* is the zero-frequency diffusivity and *R* is the coefficient of the leading frequency-dependent correction. Inserting this approximation into Eq. 7 separates the attenuation into a time-independent diffusion term and a restriction-weighted frequency-dependent term, according to

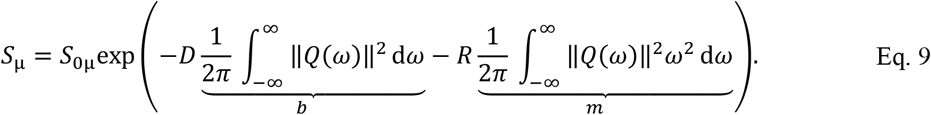

In restricted compartments with simple geometries, such as spheres or cylinders, *D*(*ω*) can be derived exactly from the size of the restriction (Stepisnik, 1993). In sufficiently simple materials, where the geometry and intrinsic diffusivity can be assumed or measured by dedicated experiments, this relation enables estimation of characteristic compartment dimensions. However, we aim to investigate heterogeneous biological tissue and therefore consider the related parameters to be phenomenological, whereby assumptions can be avoided in favor of more general, albeit less interpretable, parameters related to the diffusion process rather than specific microstructure features.

Following Lundell and Lasič (2020), we generalize Eq. 9 to a tensor-valued representation where the signal from each micro-domain can be described by domain-specific Gaussian (**D**_µ_) and restricted (**R**_µ_) diffusion tensors, according to

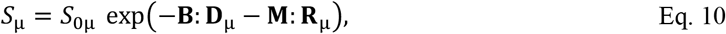

where the signal depends on both the b-tensor (Eq. 2) and the restriction weighting m-tensor

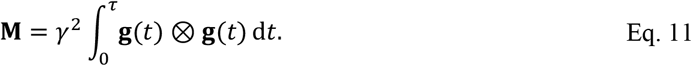

As with the b-tensor, the m-tensor has a size (*m*, Eq. 5 and Eq. 9) and a shape (ζ_m_, Eq. 6). In analogy to QTI, the powder-averaged signal in a mixture of heterogeneous micro-domains can be approximated by using another cumulant expansion (similar to Eq. 4), according to

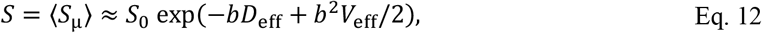

where the effective diffusivity and diffusional variance are

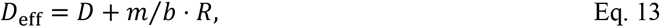

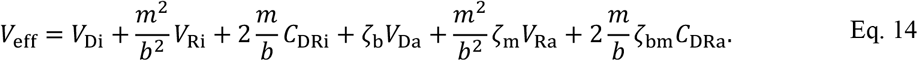

The variance terms (*V*) correspond to the variances of diffusivities, and cross-terms (*C*) are the covariances between gaussian and restricted components (Chakwizira et al., 2022, Lundell and Lasič, 2020). In this initial investigation, we assume that intra-compartment kurtosis and cross-terms are negligible (see Discussion). The unknown parameters in Eq. 13 and Eq. 14, along with their SI-units in square brackets, are related to the distributions of micro-domain tensors, according to

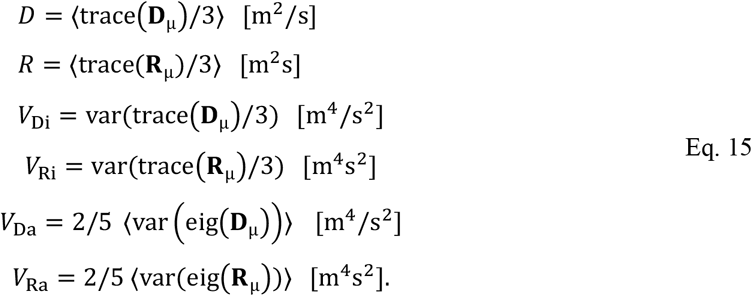

Thus, we can write Eq. 12 based on the parameters in Eq. 15 to arrive at the employed signal representation

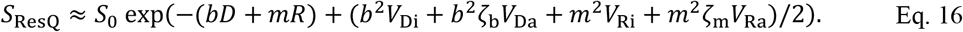

The way restricted diffusion is captured can be appreciated by considering Eq. 13 in an unrestricted, homogeneous, and isotropic medium (*V*_eff_ = 0). By factoring out the diffusion encoding strength, the logarithm of the normalized signal log(*S*/*S*_0_) = –*b*⋅*D*_eff_ = –*b*⋅(*D* + *m/b*⋅*R*) shows that the effective diffusivity is a linear function of *m/b*, such that we observe *D*_eff_ = *D* at long diffusion times (*m/b* ≈ 0) and an increasing diffusivity at shorter diffusion times (*m/b* > 0) proportional to *R*. Notably, the factor *m/b* is not a straightforward inverse of the diffusion time, but rather the variance of the encoding power spectrum. Thus, gradient waveforms that yield dephasing vectors with relatively rapid oscillations and short diffusion times will have high *m/b* and vice versa. Nilsson et al. (2017) denote this quantity *V*_ω_. We use *m*/*b* to avoid confusion with the estimated variance parameters.

It is also helpful to relate Eq. 16 to established signal representations. For clarity, we use the powder-averaged form throughout. The six ResQ parameters (*D, R, V*_Di_, *V*_Ri_, *V*_Da_, *V*_Ra_) can be estimated when the six experimental parameters (*b, b*^2^, ζ_b_, *m, m*^2^, ζ_m_) are modulated in a non-colinear fashion. If multiple b-tensor shapes are generated by spectrally matched waveforms (constant *m/b*) (Lasič et al., 2025), the representation reduces to the three unknown parameters of QTI (*D, V*_i_, and *V*_a_ in Eq. 4) in which the time-dependence of parameters cannot be decoupled. Furthermore, if a single b-tensor shape is used so that ζ_b_ is also constant, the representation reduces to diffusional kurtosis imaging (DKI) (Jensen et al., 2005), and if only low b-values are used, such that the *b*^2^-dependence is not probed, it reduces to diffusion tensor imaging (DTI) (Basser et al., 1994). Naturally, if restriction effects are negligible (all **R**_µ_ ≈ 0), QTI and ResQ are equivalent because all terms in Eq. 16 apart from *D, V*_Di_ and *V*_Da_ vanish. This is the case of ‘multi-Gaussian’ diffusion which is assumed in diffusional variance decomposition (Szczepankiewicz et al., 2016, Lasič et al., 2014) and QTI (Westin et al., 2016). To maintain a distinction between approaches that do and do not account for restricted diffusion, we will refer to these representations as ResQ (Eq. 16) and QTI (Eq. 4).

Finally, to simplify the interpretation of diffusion time-dependent parameters, and to enable a straightforward comparison to similar studies, we define parameters that capture the diffusion time dependence given the current experimental design. These parameters are denoted with a prefix ‘Δ’ to indicate a difference arising from diffusion-time dependence

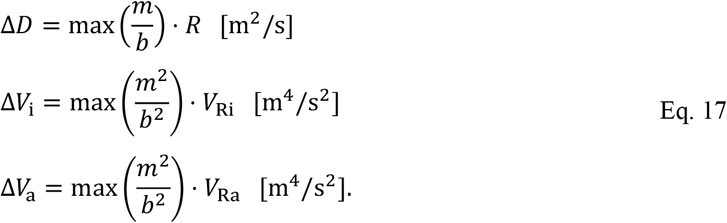

These parameters are preferred because they have the same units as their corresponding Gaussian analogues and can therefore be compared directly. The drawback is that they depend on the experiment, so conversion may be needed to compare values across studies. In this study, max(*m/b*) = 9.5·10^4^ s^−2^ such that any value of Δ*D* in units of µm^2^/ms can be converted to *R* in units of µm^2^ms by dividing the value by 9.5.

## Methods

This study comprises a longitudinal MRI investigation of mice bearing human prostate cancer xenografts. The mice were distributed into an untreated control group (CTRL) or a group receiving external beam radiotherapy (RT). All animal experiments were performed in accordance with national legislation on laboratory animal protection and permitted by the Local Ethics Committee for Animal Research at Lund University (permit number 4350-20).

### Prostate cancer tumor model

We implanted ten (*n* = 10) immunodeficient mice (BALB/c^nu/nu^, Janvier, Le Genest-Saint-Isle, France) with a human prostate cancer cell line (LNCaP, prostate carcinoma cell clone FGC ATCC®CRL-1740, lot no. 5972254) by subcutaneous injection of cells 5-7·10^6^ cells per mouse in a 200 µL cell suspension consisting of a 1:1 mixture of Matrigel (Corning) and RPMI 1640 medium. LNCaP cells have a diameter of approximately 14-21 µm when cultured and measured in vitro (Cifuentes et al., 2003). The injection was on the flank in proximity to the hind leg on the right side. The animals were regularly monitored for tumor growth by caliper and ultrasound (Roth et al., 2024), body weight, and physical signs of illness for the full duration of the study.

### Schedule for MRI and radiotherapy

Approximately 40 days after the tumor cell injection, the mice were examined by MRI following a pre-determined schedule. The first group of five mice (*n*_RT_ = 5) were treated by external radiotherapy (see below). Mice in the treatment group were examined by MRI at up to four time points, including 1-2 days before treatment, as well as 1-2 days, 1 week, and 2 weeks after treatment (*t*_RT_ ≈ [-1, 1, 7, 14] days after treatment). The second group of five untreated mice (*n*_CTRL_ = 5) was used as a control. This group was scanned up to six times with a separation of 3-10 days between each session (*t*_CTRL_ ≈ [0, 4, 8, 12, 21, 27] days after first exam). In the treatment group, two mice were terminated ahead of schedule due to issues unrelated to the treatment (injuries from fighting); one was scanned once (and was never treated), and one was scanned twice. In the untreated group, all mice were scanned four times, and one mouse was scanned six times. The total number of MRI examinations was *n*_MRI_ = 37.

The tumor volumes were estimated from the morphological MRI scans and plotted as a function of time (Figure 2). Since the tumors developed at different times, with a high inter-subject variability (Roth et al., 2024), we mapped the tumor volume trajectories onto a ‘common timeline’. This was done by fitting a quadratic reference function to the mouse with the longest inclusion period and shifting all other untreated data to best fit this reference by means of least square minimization. A single shift was used per subject, wherein mice from the treatment group only contributed their first time point to this process since it was the only point at which they were untreated.

**Figure 1.**
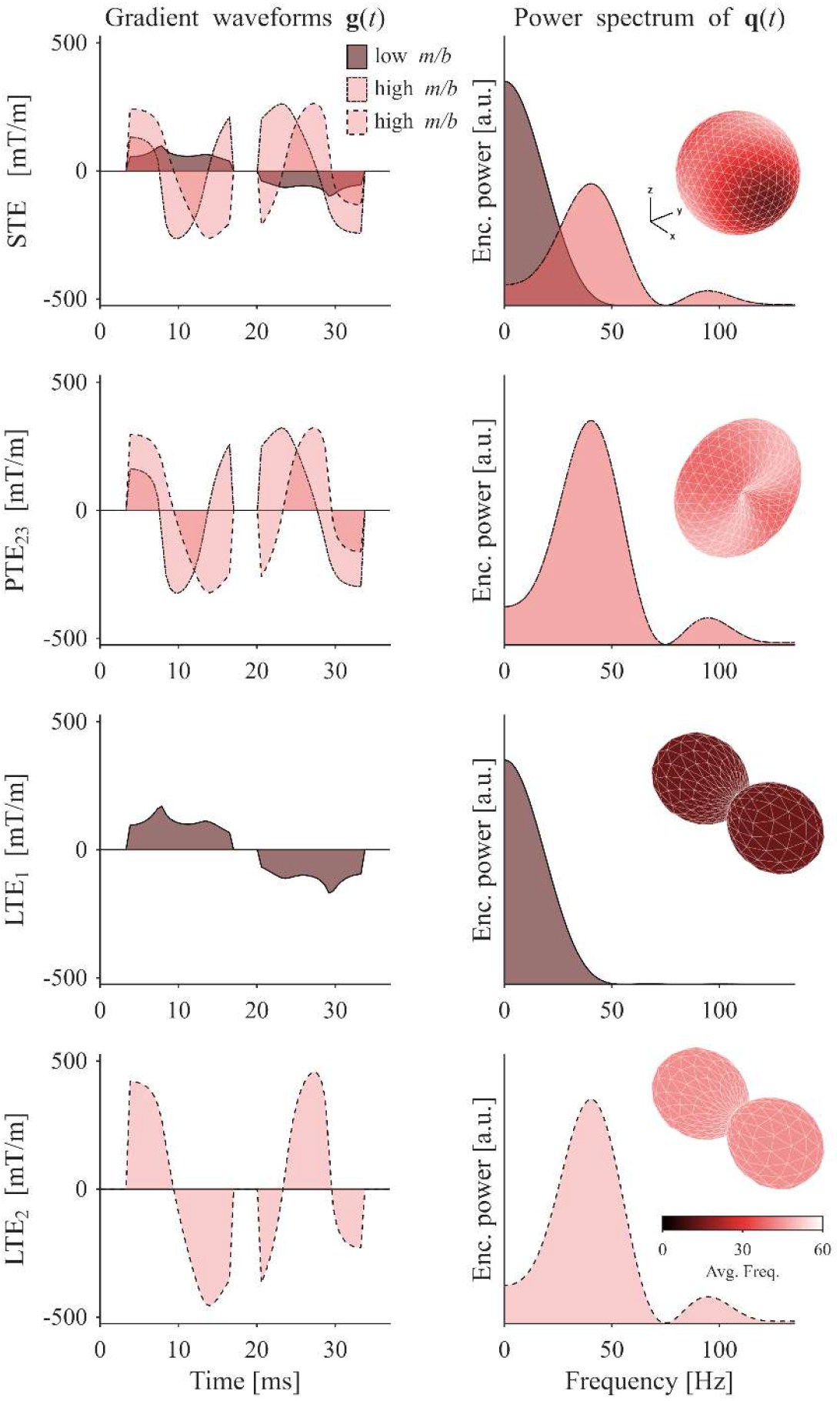
Diffusion encoding gradient waveforms used in the experiments (left column) along with the encoding power spectra (right column). The top row shows the waveform for spherical b-tensor encoding (STE), with one axis that exhibits long diffusion times (low *m*/*b*), and two axes with short diffusion times (high *m*/*b*). Thus, the STE has a relatively variable, or mixed, diffusion time characteristic along different spatial directions. The inset glyphs visualize the direction-dependent diffusion encoding and diffusion time; the distance of the surface from the center is proportional to the diffusion encoding strength and the color of the surface shows the average frequency (including only the positive side) of the encoding power spectrum. Remaining waveforms for linear and planar b-tensor encoding (LTE and PTE) were subsets of the STE, as denoted by the subscript in their names (Table 1).

**Figure 2.**
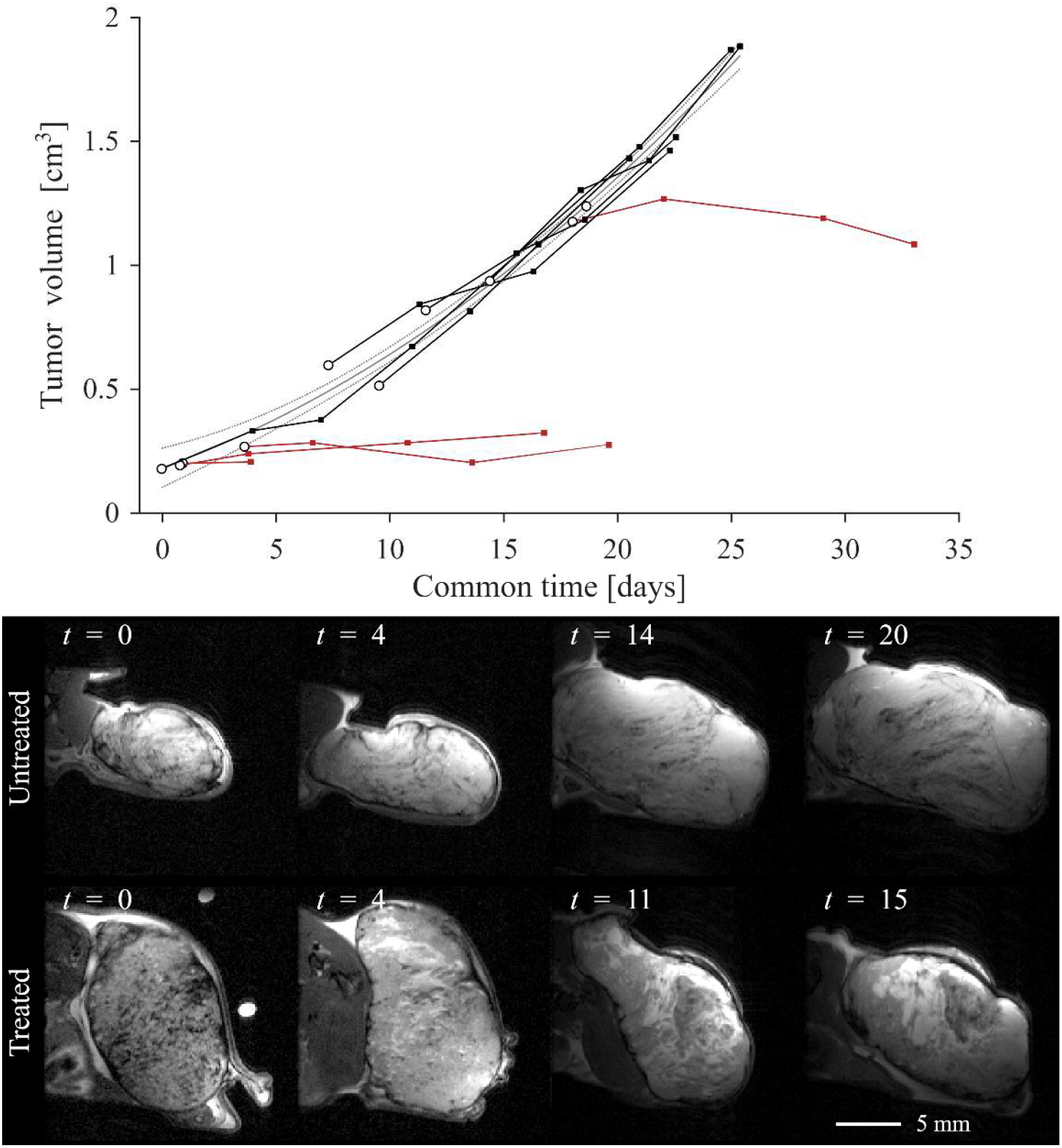
Tumor volumes mapped onto common time show that, in the absence of treatment the tumors grow rapidly (approximately 30 mm^3^/day), whereas the treatment appears to effectively suppress tumor growth. The graph shows the tumor volume as a function of common time (see Schedule for MRI and radiotherapy) where the first MRI examination is demarcated by a black circle, data from the untreated and treated groups are plotted as black and red lines, respectively. The gray lines show the quadratic trend of untreated data along with a corresponding 95% confidence interval (dashed lines). The lower panels show the morphology at four timepoints for an untreated and a treated tumor. The morphology was relatively heterogeneous between and within samples with variable structural and textural features. In the untreated example, the tumor exhibits lobular structures and a relatively variable signal intensity. By contrast, at the starting point the treated example exhibited a speckled texture, but no lobular structures. Already two days after treatment (*t* = 4 days), the morphological scan exhibits gross changes with an elevated presence of hyperintense regions indicative of necrosis and fluid buildup. At the final time (*t* = 15 days), the fluid-like necrosis is dominant, but several regions of dense tissue remain.

### Treatment by external beam irradiation

The external beam irradiation was performed at a 220 kV small-animal cabinet irradiator unit (XenX, Xstrahl Life Sciences, Atlanta, Georgia). The unit resembles a clinical linear accelerator for radiotherapy with a 360°-gantry and a collimation device enabling conformal irradiations. Each treatment was delivered using a single field tangential to the body with a conservative medial field edge to avoid unnecessary exposure of normal tissues. The absorbed dose at a reference point in the center of the tumor was 10 Gy in a single treatment fraction. The treatment was performed under isoflurane anesthesia and each procedure lasted approximately 5 minutes.

### Immunohistochemistry and structure tensor analysis

The mice were sacrificed 0-3 days after the final MRI examination. Tumors were dissected and fixed in paraformaldehyde, dehydrated, embedded in paraffin, and sent to a commercial laboratory (lmaGene-iT AB, Lund, Sweden) for immunohistochemistry and digitalization. Briefly, samples were cut into sections at a thickness of 4 µm through the bulk of the tumors. Sections were stained to show cell membranes and nuclei with hematoxylin and eosin (H&E), proliferating cells with Kiel 67 (Ki67), and prostate cancer cells with prostate specific membrane antigen (PSMA). Finally, they were scanned at a resolution of 0.2 µm. The histology was not spatially matched to the MRI data.

To quantify structural anisotropy in the histological images, we performed structure tensor analysis (Bigun, 1987) on one H&E-stained section per subject, following the methodology described by (Szczepankiewicz et al., 2016). Images were first downsampled to a spatial resolution of 1 µm and converted to grayscale by averaging the color channels. Image gradients were computed to derive the local structure tensor, and the resulting tensor field was smoothed by convolution with a Gaussian kernel with standard deviation 20 µm. Structural anisotropy was quantified as 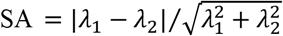 where λ_1_ and λ_2_ are the eigenvalues of the structure tensor. Structural anisotropy maps were then generated to provide a data-driven selection of regions with differing degrees of anisotropy. To reduce the influence of empty spaces on the distribution of SA, automated masking was applied. Regions representing the average and highest structural anisotropy were defined as those with SA values equal to the 50^th^ and 99^th^ percentiles of all values within the sample (see Supporting Information).

### MRI data acquisition

All mice were investigated by MRI at 9.4 T (BioSpec Avance III, ParaVision 7.0.0) with inner bore size of 12 cm, maximal gradient amplitude of 680 mT/m, a minimal rise time of 0.15 ms, a transmit-only volume coil with an inner diameter of 87 mm, and receiver coils with diameters of either 10 mm or 20 mm as best suited to the size of the tumor (Bruker, Ettlingen, Germany). During imaging, animals were anesthetized with continuous administration of isoflurane. Throughout the procedure, the breathing rate and rectal temperature were monitored using SA Instruments (Stony Brook, NY, USA) monitoring system. The temperature was maintained at approximately 37° C by circulating warm water in a heating cover.

The examination protocol included multiple scout scans, a fast spin echo morphological scan (RARE), and an advanced diffusion-weighted spin echo sequence with EPI readout that enables user-defined ‘free gradient waveforms’ (github.com/mdbudde/mcw_Preclinical_MRIsequences). The total exam time was at most 60 min. All imaging was performed in the axial plane without triggering.

The morphological RARE scan was performed with TE = 30 ms, TR = 2.5 s, FOV = 20×20×23 mm^3^, and voxel size = 125×125×1000 µm^3^, echo-train length = 8, number of averages = 6, for an acquisition time of 5:00 min.

The parameters in the advanced dMRI sequence were TE = 37 ms, TR = 2.2 s, FOV = 20×20×23 mm^3^, and voxel size = 250×250×1000 µm^3^. Four gradient waveforms were used to yield spherical, planar and linear b-tensors (see Diffusion encoding gradient waveform design), at encoding strengths of *b* = [0.1, 0.7, 1.4, 2.0] ms/µm^2^ using 30 rotations each. We also acquired 16 volumes without diffusion encoding, but these were not used in ResQ analysis to avoid signal from capillary blood. Thus, the measurement contained a total of 496 samples per voxel acquired in 18:11 min.

### Diffusion encoding gradient waveform design

We used gradient waveforms for tensor-valued diffusion encoding that were numerically optimized and compensated for concomitant gradient effects (Sjölund et al., 2015, Szczepankiewicz et al., 2019b). Additionally, we desired a greater level of control of the restriction weighting and diffusion time. Our design goals were to (*i*) maintain a high diffusion encoding efficiency (high *b* for a given encoding time) while (*ii*) making the diffusion encoding **B** and **M** tensors approximately axisymmetric. The requirement on axial symmetry means that the tensors have one or two unique eigenvalues which enables us to distribute the axis of symmetry along a given set of well-distributed directions (Bak and Nielsen, 1997, Jones et al., 1999) such that the powder average signal is rotation invariant (Szczepankiewicz et al., 2019a). By contrast, if the encoding tensors are skewed (three different eigenvalues) a more elaborate powder averaging rotation scheme must be used (Jespersen, 2025). We achieved these design goals by optimizing an axisymmetric ‘source gradient waveform’ from which all other waveforms were derived.

The duration of the source gradient waveform (**g**_source_(*t*) = [*g*_1_(*t*) *g*_2_(*t*) *g*_3_(*t*)]) was set to 13.7 ms before and after the refocusing pulse, and the separation was set to 3 ms to accommodate the refocusing pulse and accompanying slice selection gradients. To generate gradient waveforms with approximately axisymmetric encoding power spectra, we made the following modifications to the numerical optimization (Szczepankiewicz et al., 2023). First, we configured the target b-tensor to have diagonal elements [1.00 0.15 0.15]. This promotes waveforms with long diffusion times along the first axis, and short diffusion times along the second and third axes. Second, we imposed an optimization constraint on the symmetry of the gradient waveform such that the waveform on the first axis is mirrored around the central time point (*g*_1_(*t*_c_+*t*) = *g*_1_(*t*_c_–*t*)), whereas the waveform is identical along the two remaining axes but reversed in time (*g*_2_(*t*) = *g*_3_(–*t*)). Taken together, this configuration produces a waveform that yields full rank encoding tensors (rank(**B**) = 3 and rank(**M**) = 3), is approximately axisymmetric along the first axis, and maintains a high encoding efficiency.

From the source waveform, we derived a set of four gradient waveforms with complementary diffusion encoding characteristics based on the source waveform. The waveform for spherical b-tensor encoding (STE) was produced by scaling the amplitude of the source waveform (Szczepankiewicz et al., 2021), maintaining the relatively long diffusion time along the symmetry and short diffusion times in the orthogonal plane; planar b-tensor encoding was achieved by keeping only the second and third axes (PTE_23_); and the two linear b-tensors were produced by keeping only the first or second axes, which yielded linear encoding with long and short diffusion times (LTE_1_ and LTE_2_). Notably, all resulting waveforms inherit the axial symmetry in **B** and **M**, as well as the compensation for concomitant gradient effects. Note that the subscripts refer to the axes of the source waveform, but the executed waveforms were additionally rotated and scaled (Möller and Hughes, 1999).

### MRI data processing and parameter estimation

We used the native image reconstruction at the MRI system to produce magnitude images in the ParaVision format. These images were converted into 4D-nifti volumes by DSI Studio (https://dsi-studio.labsolver.org/, version: Chen, Jun 9, 2022) (Yeh, 2022). Using in-house developed code, the b- and m-tensors (Eq. 2 and Eq. 11) were calculated from the ParaVision meta data files which contain a complete record of the diffusion encoding gradient waveforms with accompanying waveform indices, amplitude scaling factors, and rotation matrices.

Two post processing steps were applied to data. First, we used MRtrix (Tournier et al., 2012) to perform denoising (Cordero-Grande et al., 2019) on the magnitude signal. Second, the data was corrected for motion and eddy-current distortions by co-registration to extrapolated image references (Nilsson et al., 2015) using ElastiX (Klein et al., 2010).Diffusion parameters were estimated voxel-wise by fitting Eq. 16 to the signal. The fitting procedure was implemented in Matlab (MathWorks Inc, MA, USA) using non-linear least-squares minimization (*lsqcurvefit*). The fitting bounds were *D* ∈ [0 3.5] 10^−9^ m^2^/s, *V*_Di_ and *V*_Da_ ∈ [–2.5 2.5] 10^−18^ m^4^/s^2^, *R* ∈ [–6 6] 10^−14^ m^2^s, and *V*_Ri_ and *V*_Ra_ ∈ [–6 6] 10^−24^ m^4^s^2^. Although negative values for the time-independent variances (*V*_Di_ and *V*_Da_) are not physical, they can be caused by noise and artifacts. We therefore allowed negative values to avoid an overrepresentation of estimates at the bounds and maintain accuracy across observations.

To estimate the corresponding parameters given the multi-Gaussian assumption, we also performed analogous estimations with QTI (Eq. *4*), i.e., under the assumption that diffusion-time effects are negligible.

To create per-tumor measurements, ROIs were manually defined to include tumor tissue. This was based on the baseline signal maps (*S*_0_) with guidance from the morphological images, while ensuring exclusion of regions with noticeable image artifacts.

Finally, to survey the data quality, we have presented single diffusion-weighted images (before/after denoising) at *b* = 0.1 and 2.0 ms/µm^2^, along with the average signal at the highest b-value in the Supporting Information. Furthermore, the signal to noise ratio at *b* = 0 ms/µm^2^ was estimated as the baseline signal from Eq. 16 divided by the estimated standard deviation of noise generated by the denoising procedure, such that SNR = S_0_/σ.

### Analysis of dMRI parameters

We performed an analysis of ResQ parameter maps and longitudinal trends across treated and untreated tumors. This included a qualitative evaluation of ResQ parameter maps that were estimated voxel-wise and shown for axial slices through the center of mass of the tumors. To highlight the ResQ parameters, we showed distributions of parameters that capture diffusion-time dependence (Δ*D*, Δ*V*_i_, and Δ*V*_a_). Furthermore, we visualized the average ResQ parameters across whole tumors as a function of common time (see Schedule for MRI and radiotherapy). To quantify the temporal trends, we reported the average ResQ parameters and their longitudinal evolution for treated and untreated groups separately. To account for treatment effects and paired longitudinal data, we used a mixed-effects model to test for linear correlations between ResQ parameters and time. This was done in Matlab using *fitlme*, wherein slopes were allowed to vary by group (*g*, treated/untreated, fixed effect *β*_1*g*_) and intercepts could vary by individual mouse (*i*, random effect *α*_0*i*_). The model was *y*_*ij*_ = *α*_0*i*_ + *β*_1*g*_ *t*_*ij*_ *+ ε*_*ij*_, where *t*_*ij*_ is the time for individual *i* at the *j*^th^ examination, and *ε*_*ij*_ is the residual error. To account for multiple testing, the significance threshold was Bonferroni-corrected to α = 0.05/12.

To investigate the impact of including or excluding the effects of time-dependent diffusion we compared ResQ vs. QTI. This was done by qualitative inspection of the signal vs. b-values in representative examples of treated and untreated tumors. The inspection of fit functions (Eq. *4* vs Eq. 16) was used to highlight how well each representation captured the dynamics of the signal. In contrast to the parameter maps, this is done for the signal averaged across the whole tumor ROI, which means that it generates a high-precision signal across a relatively heterogeneous composition of tissue. Furthermore, we quantified the difference between the analogous diffusion parameters from ResQ and QTI for a representative untreated tumor as well as from all available data. The analysis across all data compares the distribution of parameters that are estimated voxel-wise and then averaged across the tumor ROI at each exam time. Each examination therefore contributes one point of data for a total of 37 data points per parameter.

Finally, we investigated the relationship between effective diffusivity and waveform-specific restriction weighting. This was done by estimating the voxel-wise effective diffusivity, *D*_eff_, using Eq. 12 for each waveform separately (each fit has constant *m/b*, ζ_b_, and ζ_m_). This yields an estimation of *D*_eff_ that is equivalent to a DKI analysis of the powder-averaged signal, i.e., this waveform-specific diffusivity is unaffected by the relations that may otherwise be implicit from the joint analysis in Eq. 16. We could not perform the same comparison for *V*_eff_ across different waveforms as these values are conflated by the shape of b- and m-tensors. In this analysis, only used *V*_eff_ as a dummy variable that prevented higher order effects from biasing the estimation of *D*_eff_. From theory (Eq. 13), we expect the effective diffusivity to increase with *m*/*b*. Therefore, we performed linear correlation analyses on the average *D*_eff_ across tumor ROIs vs. *m*/*b* for each waveform. To quantify the linearity of the functional relationship, we report the distribution of linear correlation coefficients (Pearsson’s *r*) across all 37 MRI exams.

## Results

The set of generated gradient waveforms for tensor-valued diffusion encoding with controlled restriction weighting is presented in Figure 1 and the encoding parameters are summarized in Table 1. As intended from the optimization conditions, the STE has one axis along which the diffusion time is long, and two orthogonal axes that are identical, albeit time reversed, with short diffusion times. Despite it not being an explicit optimization target, the constituent parts of this STE waveform span nearly an order of magnitude of restriction weighting factors *m/b* between 1.17·10^4^ to 9.48·10^4^ Hz^2^ whereas the exchange weighting times, Γ as defied by Chakwizira et al. (2022), were relatively tightly grouped between 10 to 14 ms. Similarly, the PTE waveform was characterized by short diffusion times, however, the projection of the average power spectrum frequency onto the diffusion encoding glyph, reveals that there is some spectral variation even in the PTE. This means that the encoding spectra appear identical in the rotation shown in Figure 1, but the interactions between axes yield a slight deviation from axial symmetry. Finally, the two LTE waveforms have identical spectra as the short and long diffusion time axes of the STE, respectively. For reference, the highest gradient amplitude that was required to achieve a b-value of 2.0 ms/µm^2^ was 457 mT/m.

**Table 1.** The set of four diffusion encoding gradient waveforms were derived from the source waveform as defined in the construction column. Note that the waveforms were also scaled to achieve the sought b-tensor shapes and b-values (Szczepankiewicz et al., 2021), as well as rotated to achieve a distribution of samples that yielded an accurate powder average.

| Name | Construction | b-value<br>$b$ [ms/$\mu\text{m}^2$] | Restr. weight<br>$m/b$ [s$^{-2}$] | b-shape<br>$\zeta_b$ [1] | m-shape<br>$\zeta_m$ [1] |
| --- | --- | --- | --- | --- | --- |
| STE | $[g_1(t) \ g_2(t) \ g_3(t)]$ | 0.1, 0.7, 1.4, 2.0 | $6.71 \cdot 10^4$ | 0.00 | 0.20 |
| PTE<sub>23</sub> | $[ \ 0 \ g_2(t) \ g_3(t)]$ | | $9.48 \cdot 10^4$ | 0.25 | 0.28 |
| LTE<sub>1</sub> | $[g_1(t) \ 0 \ 0 \ ]$ | | $1.17 \cdot 10^4$ | 1.00 | 1.00 |
| LTE<sub>2</sub> | $[g_2(t) \ 0 \ 0 \ ]$ | | $9.48 \cdot 10^4$ | 1.00 | 1.00 |

The longitudinal tumor volumes in Figure 2 show that radiotherapy effectively inhibited tumor growth. Corresponding morphological MRI images further show a progressive loss of viable tissue in the treated tumor. A qualitative evaluation of tumor histology revealed immunohistochemical differences between treated and untreated tumors. As shown in Figure 3, all tested stains show marked changes, with the most prominent differences observed in proliferative activity (Ki67) and expression of prostate specific membrane antigen (PSMA) where treated tumors displayed more heterogeneous and lower levels of positive staining. Moreover, histology revealed that tumor tissue mostly appeared isotropic, lacking prevalence of structures that would predict elevated diffusion anisotropy (see Supporting Information).

**Figure 3.**
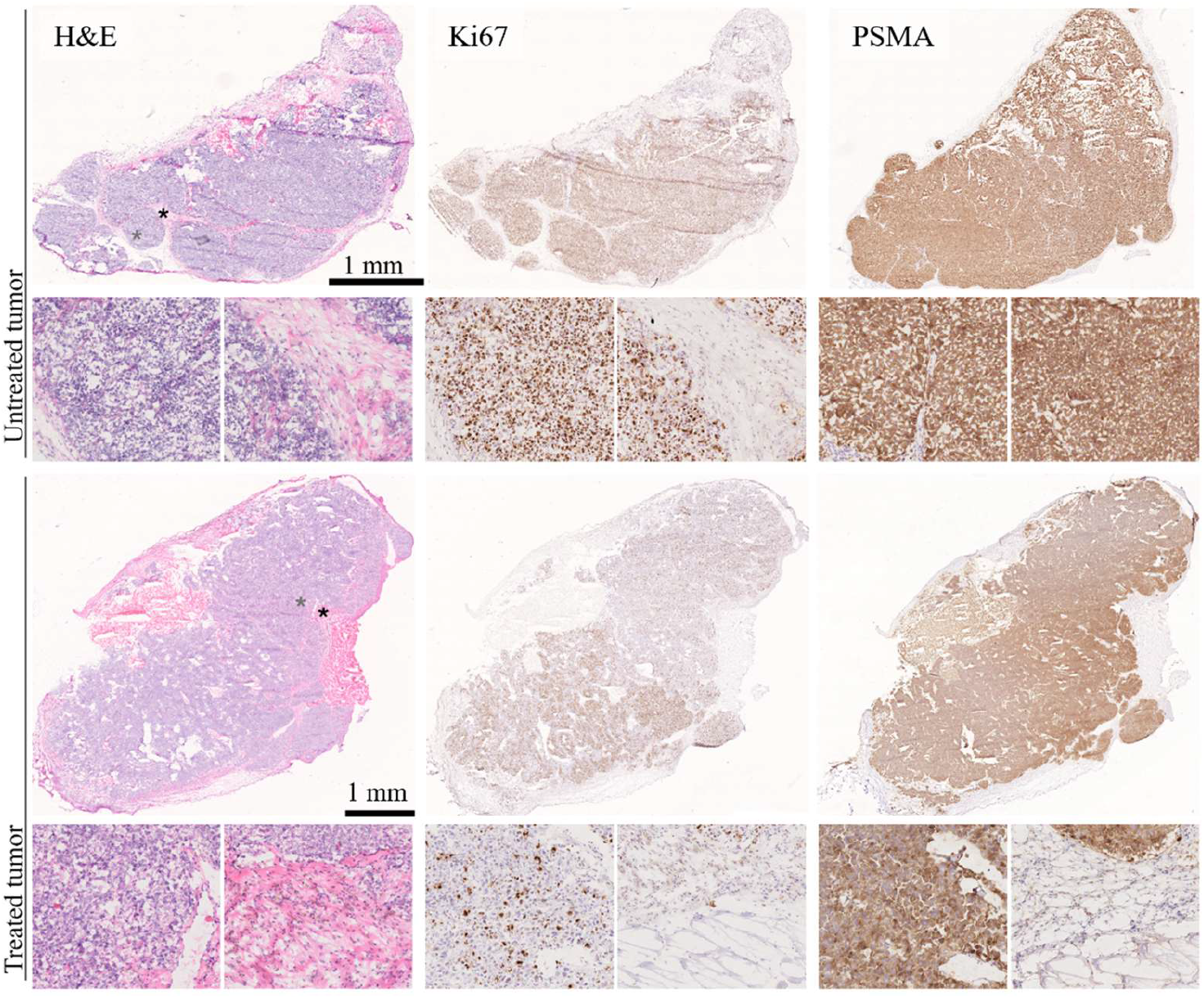
Histology and immunohistochemical staining of tissue sections for representative cases of treated or untreated tumors (same tumors as in Figure 2). The large images show whole sections through the solid parts of the tumors. Pairs of magnified H&E images show regions where structural anisotropy was equal to the 50^th^ (left, gray asterisk) and 99^th^ percentiles (right, black asterisk), i.e., examples of normative and relatively high structural anisotropy in each tumor. Magnifications of remaining stains are from approximately the same regions. The H&E images indicate that tumors were not homogeneous but grew in sub-lobules separated by stromal tissue. The Ki67 (a marker for cell proliferation) showed a marked difference between the treated and untreated tumors. The untreated tumor was homogeneously positive for Ki67, whereas the treated tumor was less positive overall, and had highly heterogeneous staining within the tumor, with regions of positive clusters and large regions that were sparsely positive. This indicates that the treatment had an effect, but that the effect may have been different across different parts of the tumor. The PSMA stain was relatively homogeneous and positive throughout both tumors, albeit showing a higher variation in the treated case. The magnifications show that structural anisotropy was low throughout, where high values were rare and found in patches of stroma, or in the transition between tissues. Note that the width of magnified image is approximately 300 µm, and that similar plots for the other tumors can be found in Supporting Information.

Overall, the quality of dMRI data was fair but had a wide distribution of tumor-averaged SNR values, such that the 1^st^, 2^nd^, and 3^rd^ quartiles were 11, 14, and 22 across all sets of data at *b* = 0 ms/µm^2^. We have provided a closer inspection of data quality, and the effect of denoising, for the sample that was closest to the global average SNR in the Supporting Information.

Parameter maps from ResQ in treated and untreated tumors show a large difference between the two groups (Figure 4). The heterogeneous nature of tumor morphology, as seen in Figure 2, is also reflected in the diffusion parameter maps. The parameter maps are relatively homogeneous in the untreated case, whereas in the treated case, they exhibit high intratumor heterogeneity where both diffusivity (*D*) and isotropic diffusional variance (*V*_Di_) indicated gross differences between the medial and lateral regions of the tumor. Since the line between the two regions appears approximately tangential to the body, these differences may be due to an uneven or incomplete coverage of radiotherapy. The anisotropic variance (*V*_Da_) was low for both cases apart from a limited region in the lateral part of the treated tumor. The quality of parameter maps was generally good, although comparison with the morphological images revealed several regions with marked geometric distortions due to local field inhomogeneity. Importantly, there is a visible dependency of diffusivity on the restriction weighting of the gradient waveforms as the diffusivity at high *m/b* is markedly higher than at *m/b* = 0. The same trend, although not as prominent, was seen for the isotropic variance.

**Figure 4.**
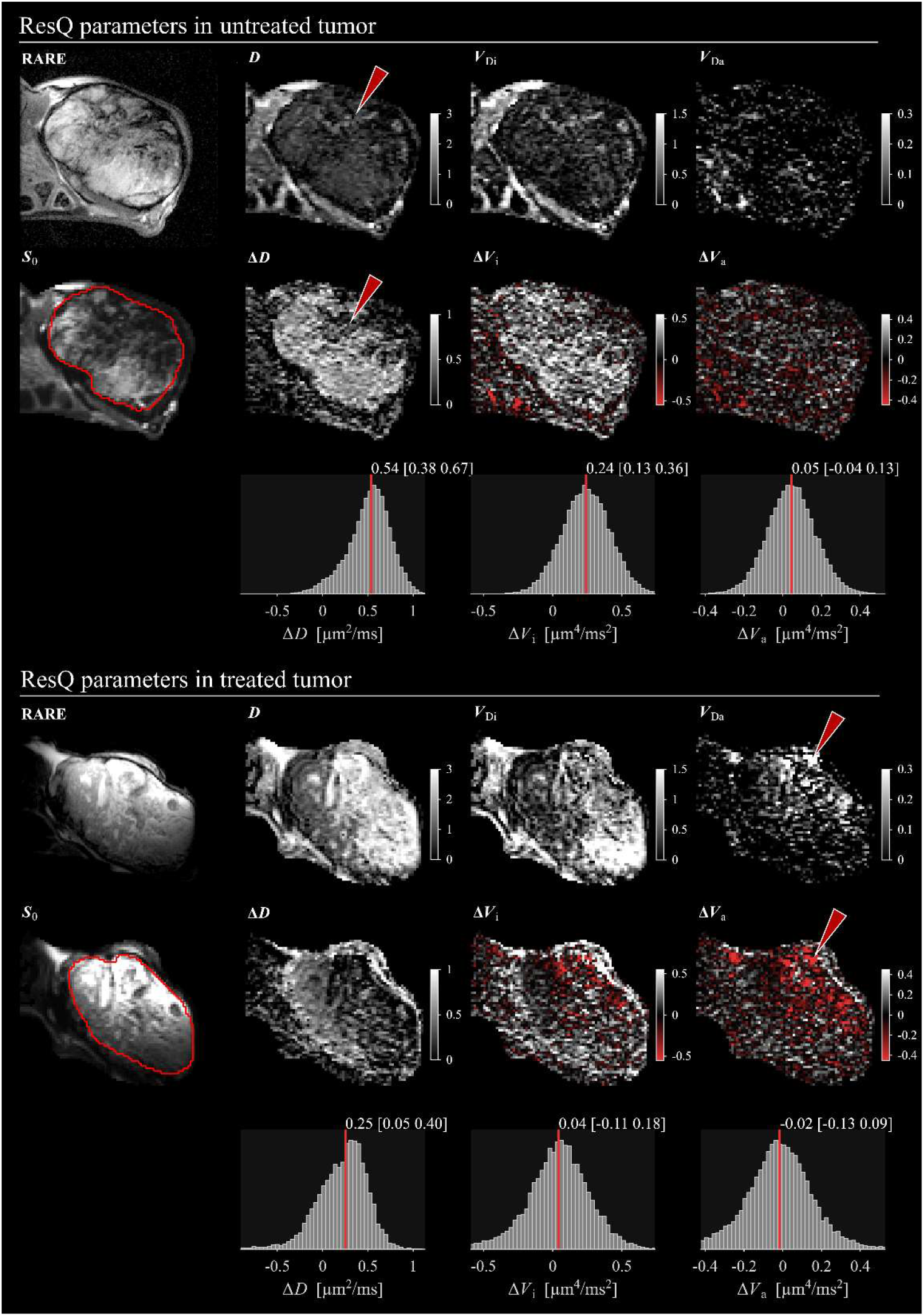
ResQ parameter maps in examples of untreated and treated mice from the final MRI exam. In the untreated tumor (top), the parameter maps were relatively homogeneous except for a region of elevated diffusivity and isotropic variance, as well as reduced restriction effects (arrows), indicating heterogeneous tissue. The treated case (bottom) exhibited prominent intra-tumor heterogeneity across all parameters. We note that the two distinct regions in the parameter maps could be caused by a radiation field that was tangential to the body but did not irradiate the medial tissue (left side of image). This case also had a region of slightly elevated anisotropic variance (arrows), although it is relatively low compared to the isotropic variance (compare color scales). In both cases, the histograms show the distribution of time-dependence parameters Δ*D*, Δ*V*_i_, and Δ*V*_a_ from within the whole tumor ROI; the red line shows the median, and the associated numbers report the median and IQR. The effect of diffusion time was visible in the diffusivity, where Δ*D* was positive. The untreated case also showed a small but positive Δ*V*_i_, which the treated case did not. The treated case also had a region of elevated *V*_Da_ and negative Δ*V*_a_ (orange arrows), indicating the presence of tissue that causes anisotropic diffusion which is reduced as the diffusion time is reduced. Units for the parameter maps are the same as stated in the histograms.

ResQ parameters were also investigated longitudinally (Figure 5). As for the example cases in Figure 4, the most prominent trend was that radiotherapy caused *D* and *V*_Di_ to increase, and Δ*D* to decrease, immediately after the radiotherapy. The mixed-effects model showed that these three parameters also changed significantly over time (Table 2) in both treated and untreated mice. However, the rate of change in the parameters was markedly different between the groups. Most notably, the diffusivity and isotropic Gaussian heterogeneity, *D* and *V*_Di_, increased in both groups, but did so at markedly higher rates in the treated group.

**Table 2.** Group-average ResQ parameters for untreated and treated tumors. In each cell we report the group average parameter followed by the expected change in 30 days as well as the standard error of this estimate in parentheses. The rate of change is based on the mixed effects correlation analysis, and an asterisk signifies statistical significance (*p* < 0.05/12).

| | $D$<br>[$\mu\text{m}^2/\text{ms}$] | $\Delta D$<br>[$\mu\text{m}^2/\text{ms}$] | $V_{Di}$<br>[$\mu\text{m}^4/\text{ms}^2$] | $\Delta V_i$<br>[$\mu\text{m}^4/\text{ms}^2$] | $V_{Da}$<br>[$\mu\text{m}^4/\text{ms}^2$] | $\Delta V_a$<br>[$\mu\text{m}^4/\text{ms}^2$] |
| --- | --- | --- | --- | --- | --- | --- |
| Untreated | 0.68 | 0.56 | 0.14 | 0.15 | 0.03 | 0.01 |
|  | +0.25 (0.06)* | -0.22 (0.03)* | +0.25 (0.02)* | -0.02 (0.05) | +0.03 (0.01) | +0.01 (0.01) |
| Treated | 1.02 | 0.51 | 0.30 | 0.15 | 0.02 | 0.01 |
|  | +1.59 (0.25)* | -0.31 (0.08)* | +0.43 (0.08)* | -0.16 (0.09) | +0.03 (0.02) | +0.01 (0.01) |

**Figure 5.**
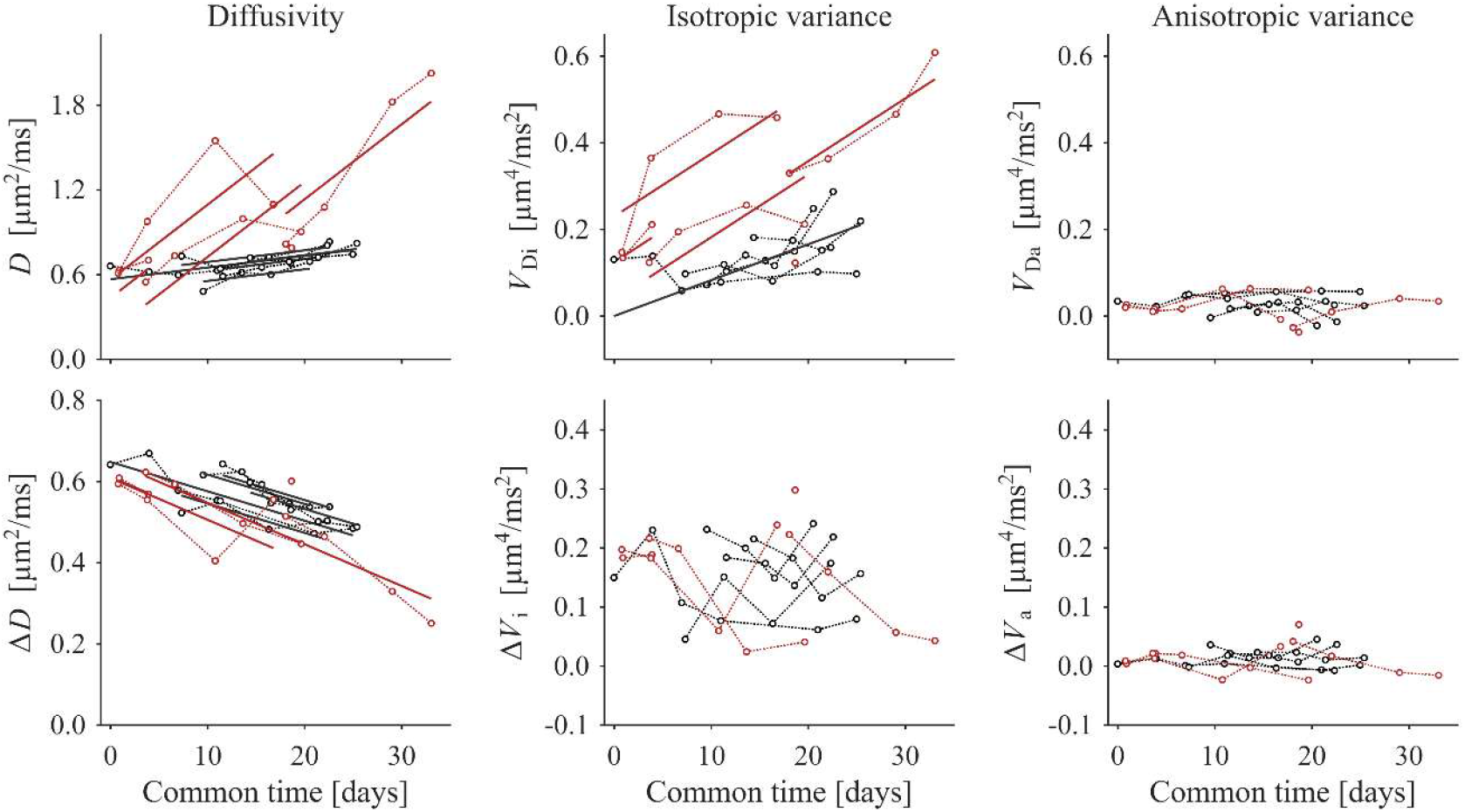
Longitudinal view of ResQ parameters with trend lines (solid lines) for parameters where these were statistically significant. The parameters are calculated as the mean across the whole tumor ROIs and plotted against the same common time as in Figure 2. Each individual subject contributes several data points (circles) connected by a dotted line (black/red are untreated/treated). The parameters most obviously affected by radiotherapy are the diffusivity and isotropic diffusional variance (*D* and *V*_Di_), which are both markedly elevated (Table 2). Furthermore, positive trends can be seen for *D* and *V*_Di_, whereas the trend in Δ*D* is negative. The diffusion anisotropy parameters (*V*_Da_ and Δ*V*_a_) are close to zero and do not correlate significantly with time.

The investigation of signal vs. b-values along with fitting functions (Figure 6) revealed that ResQ was able to capture the dynamics of the signal, whereas QTI exhibited large residuals. Indeed, the effect of diffusion time was clearly visible in the markedly different signal slopes at low b-values as measured by different waveforms. Furthermore, Figure 6 shows that the increased diffusivity caused by treatment resulted in measurements at lower signal levels. Thus, measurements in treated tumors may be more exposed to noise floor effects.

**Figure 6.**
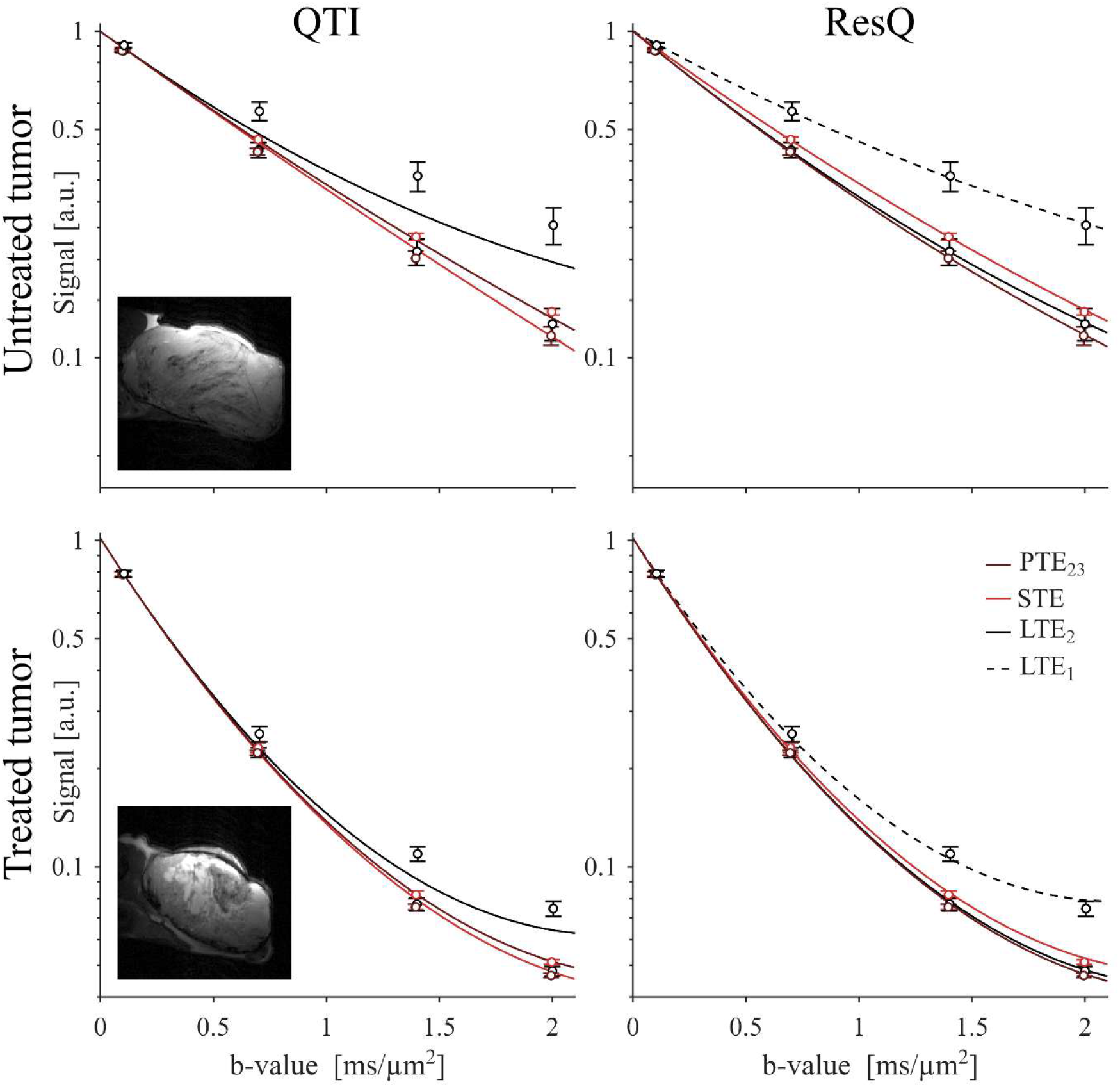
Signal vs. b-values along with fits of the representations for QTI (left column) and ResQ (right column) in untreated and treated tumors at the time of the final MRI exam (same tumors as in Figure 2). Markers show the average signal across the whole tumor volume and the whiskers show two standard deviations of the per-shot signal average. The left column shows that QTI is unable to accurately capture the dynamics of signal (lines do not coincide well with data points) that is diffusion-weighted with waveforms that provoke variable restriction effects. Indeed, the initial slope of the signal vs b-value is clearly not equal across waveforms. By contrast, the right column shows that ResQ can capture these dynamics and the lines show only small deviations from the average signals. Notably, signal averaging across the whole tumor ROI may disguise potential regions where the signal is markedly affected by the rectified noise floor. This is especially true in the treated case, where the dynamic range of signals is larger due to the relatively high diffusivity.

A comparison of QTI and ResQ parameters in an untreated case shows large differences in the estimated parameters (Figure 7). The parameter maps are calculated from the same data but exhibit large differences due to time-dependent diffusion. The most prominent error appears in the anisotropic diffusion variance, which is highly inflated in QTI analysis, whereas the isotropic variances were similar across methods.

**Figure 7.**
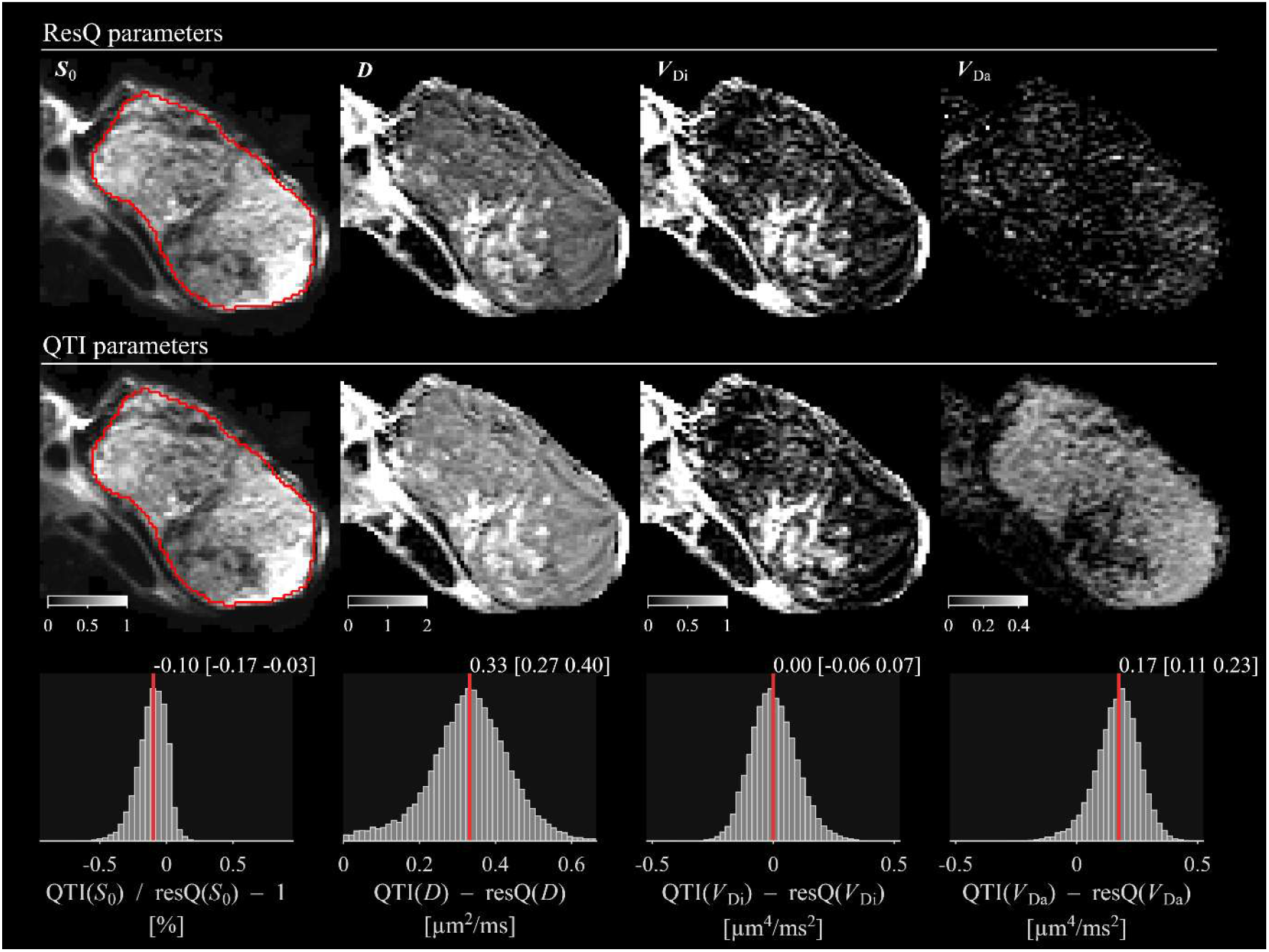
Comparison of parameter maps from ResQ and QTI. The most prominent differences can be seen in the diffusivity and anisotropic diffusional variance (*D* and *V*_Da_) which are both markedly inflated by QTI. The bottom row shows the distribution of per-voxel parameter differences across the whole tumor, the red line marks the median and the numerical values show the median and the IQR. This example highlights the potential for gros bias caused by neglecting diffusion-time effects when such are highly relevant. Note that the units for the color bars are the same as stated in the histograms.

The same trends were seen across all examinations (Figure 8). The distribution of parameters shows that the largest differences between analogous parameters derived from QTI and ResQ are the diffusivity and anisotropy parameters. It is expected that QTI would report an intermediate diffusivity as it is determined by a weighted average across diffusion encoding waveforms, whereas ResQ estimates diffusivity at *m/b* = 0 as well as its dependence on *m/b*, i.e., both *D* and *R*. However, the same is not true for the anisotropic diffusional variance, which is reported to be close to zero by ResQ but is grossly overestimated by QTI. This can be explained by the fact that the signal difference across waveforms is interpreted as microscopic anisotropy rather than different effective diffusivities across waveforms (Szczepankiewicz et al., 2016, Lundell et al., 2019, Lasič et al., 2025). Finally, the relation between effective diffusivity and the restriction weighting appeared remarkably linear (Figure 9). Indeed, all correlation coefficients were above 0.99, indicating that a linear model explains the functional relation almost perfectly (median coefficient of determination was 0.999).

**Figure 8.**
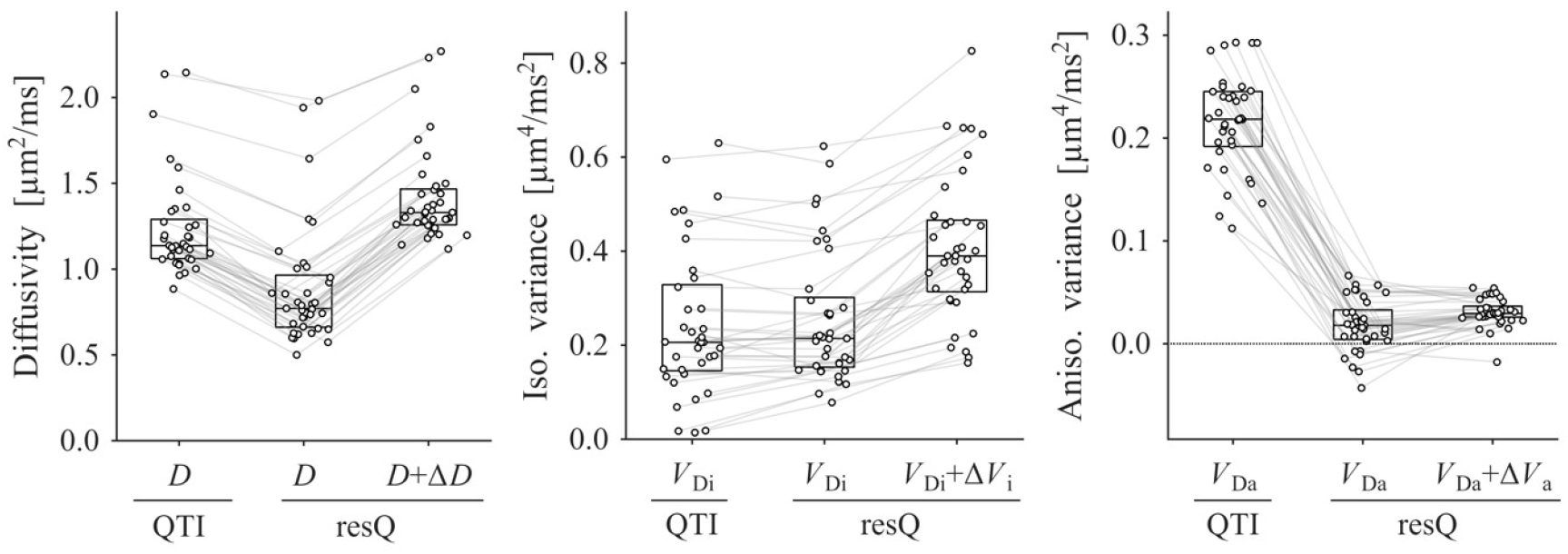
Comparison of parameters from QTI and ResQ show that restriction effects manifests consistently in all three diffusion parameters and across all examinations. Most notably, the anisotropic variance is grossly overestimated by QTI (right plot). In the absence of time-dependent diffusion, the parameters would fall on a horizontal line. Each circle is the voxel-wise parameter estimations averaged across the whole tumor ROI for each one of the *n*_MRI_ = 37 examinations. Lines connect data points from the same examination. Box plots show the median and IQR.

**Figure 9.**
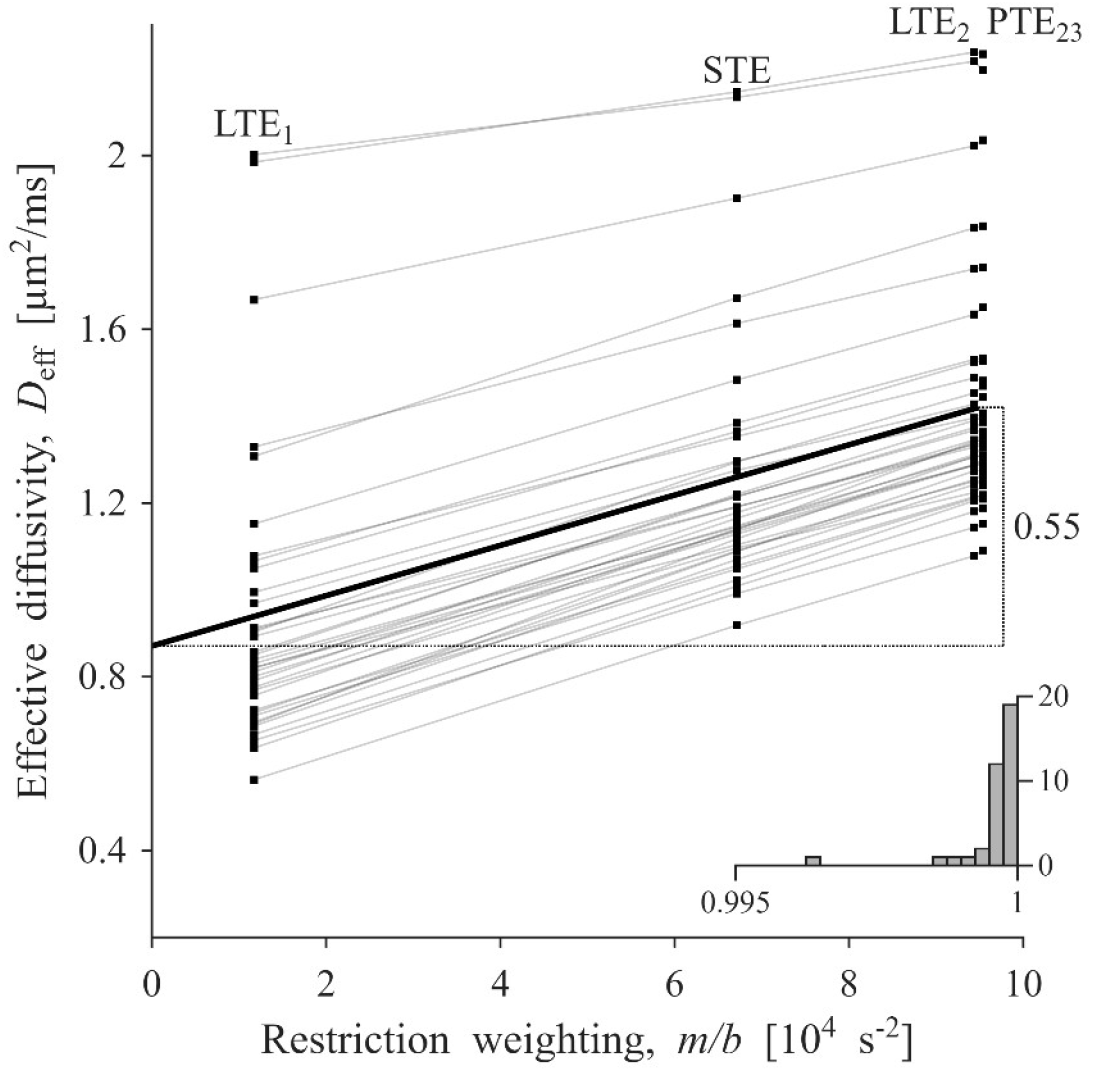
Effective diffusivity (*D*_eff_) as a function of restriction weighting (*m/b*) appears strikingly linear, as expected from theory (Eq. 13). Each data point is the average *D*_eff_ across the whole tumor ROI, and the data from each of the *n*_MRI_ = 37 examinations are linked by lines. Note that LTE_2_ and PTE_23_ have identical *m/b*, but that they are graphically separated to improve legibility. The black line shows the average linear trend, which indicates that the diffusivity increases by 0.55 µm^2^/ms as the restriction weighting goes from 0 to 9.5⋅10^4^ s^−2^. The inset histogram shows the distribution of linear correlation coefficients, where all values were above 0.995, indicating a near-perfect linear relationship between *D*_eff_ and *m/b*.

## Discussion

We have presented an initial investigation of prostate cancer xenografts by a novel framework for measuring and quantifying diffusion parameters that combines tensor-valued diffusion and restriction encoding. We demonstrated that ResQ could effectively capture the relevant signal dynamics and their dependency on restriction weighting. The parameter maps were generally of good quality and exhibited contrasts that were plausible in relation to morphological MRI and histological findings. The intra-tumor variation was especially pronounced after radiotherapy (Figure 4). A radiotherapy-induced increase in diffusivity was first reported in human prostate cancer patients by Foltz et al. (2013), and several other studies have corroborated this finding on both short and long time scales (Song et al., 2010, Almansour et al., 2023, Pasquier et al., 2018). Despite the relatively small group sizes, we could establish longitudinal trends of ResQ parameters, indicating that the onset of radiotherapy effects are rapid (Figure 5) and that these parameters distinguish treated from untreated tissue. Taken together, these findings suggest that the parameters are sensitive to both subtle and gross microstructural changes as tumors develop and are perturbed by therapeutic intervention.

We emphasize that the observed interplay between gradient waveforms and time-dependent diffusion was not a subtle effect but could be clearly seen in the diffusion-weighted signal across waveforms with different restriction weighting. Indeed, the value of diffusivity and its time-dependence, *D* and Δ*D*, were comparable in magnitude, meaning that the effective diffusivity within a measurement varied by approximately 0.5 μm^2^/ms or 60% of the minimal value (Table 2 and Figure 9). These results align with similar studies that use oscillating gradient waveforms purposefully designed to probe a wide range of diffusion times (Jiang et al., 2023, Baron and Beaulieu, 2014, Does et al., 2003), and serve as a complement to investigations with pulsed field gradients at intermediate-to-long diffusion times in prostate cancer (Lemberskiy et al., 2017).

In addition to the waveform-dependent diffusivity, the ResQ analysis revealed that the dominant higher-order effect was the isotropic diffusional variance, which reflects tissue heterogeneity, as opposed to a consistently small contribution from the anisotropic diffusional variance, which reflects microscopic anisotropy (Szczepankiewicz et al., 2016). The impact of diffusion time was most prominent in the diffusivity (Δ*D*) and isotropic diffusional variance (Δ*V*_i_), but quite weak for the anisotropic diffusional variance (Δ*V*_a_) (Figure 7), possibly due to the generally low levels of micro-anisotropy in the investigated tissue (Figure 3).

Surprisingly, we observed that isotropic diffusional variance increased with *m/b* (Δ*V*_i_ > 0, Figure 8). Given that shorter encoding times should render the diffusion process less sensitive to restrictions and therefore less revealing of tissue heterogeneity, we expected Δ*V*_i_ to be negative, although positive values are physically allowed. For example, a positive Δ*V*_i_ is expected in a mixture of Gaussian and restricted compartments that have similar effective diffusivity (*D*_eff_) at low *m/b* but becomes more heterogeneous at higher *m/b* as *D*_eff_ in the restricted compartment increases. An alternative explanation is that an interaction between the effective diffusivity and the rectified noise floor causes an *m/b*-dependent signal bias (Gudbjartsson and Patz, 1995), i.e., waveforms with high *m/b* tend to exhibit faster signal decay and therefore suffer a lower signal-to-noise ratio. This confounder is known to affect the isotropic and anisotropic variance to different degrees (Szczepankiewicz et al., 2019a) and should be taken into account when analyzing diffusion parameters.

As predicted by theory, we found that the relationship between the effective diffusivity across the four diffusion encoding waveforms was strikingly linear as a function of the restriction weighting factor, *m/b*. Since all linear correlation coefficients were tightly grouped around unity (Figure 9), we conclude that their relationship, at least in the probed interval of *m/b*, is linear and conforms to the expected behavior described by Eq. 13. We emphasize that this cannot be a side effect of the implicit constraints of the ResQ representation (Eq. 16) since the analysis considered each waveform separately, eliminating any crosstalk. This observation is promising for the utility of ResQ but does not rule out other functional forms for the time-dependent diffusivity (Eq. 8). Therefore, further study is warranted to also investigate the relationships in other tissue types, wider intervals of *m/b*, and for higher-order contributions.

By comparing ResQ to QTI, we demonstrated the potential pitfalls of ignoring diffusion-time effects when quantifying and interpreting diffusion parameters. Although the parameter maps from QTI appear feasible, closer inspection of the signal and fitting functions revealed that it could not capture the dynamics of the diffusion-weighted signal. For example, a consequence of the multi-Gaussian assumption is that initial slopes are identical across waveforms which means that the separation of signals acquired with different b-tensor shapes is attributed to microscopic anisotropy (Szczepankiewicz et al., 2016), but the separation can also be caused by varying initial slopes due to restrictions and time-dependent diffusion (Szczepankiewicz et al., 2021, Lundell et al., 2019, Lasič et al., 2025, Jespersen et al., 2017). We have demonstrated that this mechanism can cause a gross overestimation of parameters related to microscopic anisotropy (Figure 7 and Figure 8), which is corroborated by the corresponding histology which indicated limited or negligible presence of anisotropic structures. Thus, despite not having access to the ground truth, we could show that the multi-Gaussian assumption fails in prostate cancer with the likely consequence of impeded parameter accuracy and interpretation. Although the exact impact depends on the experimental setup and targeted tissue, the bias is likely to appear whenever the encoding times are commensurate with the sizes of the restriction, i.e., for short encoding times and/or large restriction sizes. This should motivate users of similar methods to investigate diffusion time characteristics as a potential confounder or leverage time-dependent diffusion to use it as a biomarker.

The strong diffusion-time effects observed in this study are particularly noteworthy because the gradient waveforms were not explicitly designed to provoke such effects. Rather, the waveforms were optimized to maximize encoding efficiency, axial symmetry, and negligible concomitant gradient effects (Sjölund et al., 2015, Szczepankiewicz et al., 2019b) to facilitate accurate measurements and straightforward translation to human MRI systems. Despite this, the nature of highly efficient non-linear b-tensors promotes dephasing vectors that include a variety of diffusion encoding times, or restriction weights (*m/b*) (Szczepankiewicz et al., 2021). This highlights the unintended, and often overlooked (Jespersen et al., 2018, de Swiet and Mitra, 1996), consequence of using combinations of gradient waveform designs without accounting for their less obvious differences (Szczepankiewicz et al., 2021).

An alternative strategy is to avoid the effects of time-dependent diffusion by gradient waveform design. Lasič et al. (2025) recently proposed two strategies to achieve this. One involves tuning the gradient waveforms such that STE and LTE sense the same apparent mean diffusivity given a specific diffusion spectrum. Alternatively, a geometric signal average of orthogonal LTE projections can be used to harmonize restriction effects across gradient waveforms. These approaches allow a simpler signal representation and facilitate a more accurate estimation of microscopic anisotropy. In parallel, Lasič et al. (2025) proposed the ‘spectral principal axis system’ which can be used to maximize sensitivity to restricted diffusion to provide additional time-dependent diffusion contrast, similar to ResQ. In lieu of waveforms with well-controlled restriction weighting or tuning, we recommend users of similar methods to investigate the per-waveform effective diffusivity, as in Figure 9, to discern if their specific combination of experiment and subject is affected by time-dependent diffusion.

Importantly, the resulting ResQ parameters must be interpreted within an appropriate context, i.e., the pre-clinical MRI setting, which enables strong gradients and relatively short diffusion times, as well as the specific subject of LNCaP prostate cancer xenografts, which exhibit highly heterogeneous microstructure. By contrast, a similar implementation for studying human prostates in vivo will likely use weaker gradients with longer diffusion encoding times to achieve similar b-values, thereby probing longer diffusion times with less prominent diffusion-time effects (Lemberskiy et al., 2017). Furthermore, we anticipate significant microstructural differences between prostate cancer xenografts grown in mice and those found in human patients. For example, our xenografts exhibited negligible-to-low microscopic diffusion anisotropy, whereas human prostate tissue is known to contain, for example, stromal tissue that exhibits anisotropic diffusion (Bourne et al., 2012). Non-negligible microscopic diffusion anisotropy has been reported in two preliminary studies of human prostate cancers (Langbein et al., 2021, Nilsson et al., 2021), however, these studies did not account for diffusion-time effects and may therefore be subject to the bias from time-dependent diffusion, as illustrated in Figure 6. Nevertheless, the introduction of high-performance human MRI systems capable of prostate imaging, both pre-clinical (Molendowska et al., 2022) and clinical (Zhu et al., 2024, Bischoff et al., 2024, Molendowska et al., 2025b), can probe relatively short diffusion times in humans. Indeed, we have demonstrated a similar diffusion-time dependence in a preliminary study of patients on a system with 300-mT/m gradients (Molendowska et al., 2024).

To our knowledge, this is the first study that reports ResQ parameters, as well as restriction-weighted tensor-valued diffusion encoding in prostate cancer. Closely related parameters can also be derived from non-parametric representations based on inverse Laplace transform in the context of ‘highly multidimensional’ measurements, such as those recently demonstrated in studies of brain (Johnson et al., 2024, Narvaez et al., 2022). Given an accurate inverse Laplace transform (Epstein and Schotland, 2008, Ronen et al., 2006), the non-parametric approach enables investigation of a wide range of parameters and parameter correlations. Although it may require larger datasets to enable accurate reconstruction of intra-voxel parameter distributions, we expect that our data could support such an analysis. However, this was outside the scope of the current study.

This study had several limitations. The main limitations originate from the assumptions of the resQ representation, where each is a potential source of bias and misinterpretation. First, we use *m/b* and *R* as convenient parameters to describe the restriction weighting of arbitrary gradient waveforms and the corresponding restriction parameter. This approach is based on the low frequency approximation which is valid when diffusion times are relatively long compared to the sizes of restrictions (Lundell and Lasič, 2020, Stepisnik, 1999, Nilsson et al., 2017). Thus, it has a limited interval of validity and may become inaccurate at high *m/b* or for large restriction sizes. The representation also does not account for specific restriction geometries or for extracellular spaces where the diffusion spectra may follow different forms, including fractional power laws (Novikov et al., 2014, Novikov et al., 2011). In heterogeneous tissue, these different spectral contributions are unlikely to be uniquely distinguishable with the present parameterization. If the true spectrum over the sampled frequency range is not well described by the quadratic approximation, the fitted *R* and corresponding variance terms become effective descriptors whose values may depend on the waveform set. Taken together, the estimated parameters capture restriction effects but cannot be straightforwardly converted to estimates of the underlying compartment geometry or size. As such, ResQ is a step toward improving methods like QTI, which may suffer from ignoring diffusion-time effects entirely. Future studies should investigate how well the assumptions hold across different tissues and how the geometry and composition of restrictions impact the interpretability of estimated parameters. Second, we use the Gaussian phase approximation, which is valid at low diffusion encoding (Stepisnik, 1999, Topgaard, 2025) but could be violated in the presence of exchange or intra-compartmental kurtosis. If omitted, these effects may bias the apparent variance terms estimated by ResQ. Related effects have been studied recently by ‘correlation tensor imaging’ (CTI), and several studies have demonstrated cases where intra-compartmental kurtosis is not negligible (Novello et al., 2022, Henriques et al., 2020, Alves et al., 2022, Chakwizira et al., 2025). In a study of mouse brains, Henriques et al. (2021) showed that the presence of microscopic kurtosis can introduce significant bias in both isotropic and anisotropic diffusional variance parameters. As CTI generally employs waveforms that are matched with respect to *m/b*, it probes features different from those of ResQ, suggesting that the methods may provide complementary information (Chakwizira et al., 2025) with potential benefits to investigations of cancers. Third, the current implementation of ResQ assumes that the covariance terms between Gaussian and restricted contributions are negligible. This assumption was first motivated by simplicity (Szczepankiewicz et al., 2023) and imposed by the current experimental design which does not support the estimation of the cross-terms. Although, to the best of our knowledge, no evidence has yet been presented that either supports or refutes this assumption, we have recently reported preliminary results indicating that covariance terms are a relevant contribution but require an extended experimental design (Molendowska et al., 2025a). If this is the case, we expect the ignored cross-terms to bias remaining parameters. Fourth, ResQ assumes that the compartments within a given voxel are non-exchanging, that is, water molecules tend to remain within a single domain for the duration of each diffusion encoding event. Chakwizira et al. (2022) recently proposed a framework that unifies both restriction and exchange effects for linear diffusion encoding, which may enable more accurate analysis in the case that both exchange and restrictions have a relevant effect in prostate tissue. Based on the present study, the effects of restrictions appear to be dominant, as supported by the large difference between signals at low b-values, and as expected for the small difference in exchange weighting times across waveform designs (10 to 14 ms). Finally, our infrastructure did not allow light microscopy to be spatially registered to the same coordinate system as the MRI data, preventing a spatially matched comparison of diffusion parameters to analogues derived from histology. This objective will be pursued in future studies to establish an accurate interpretation of relevant parameters by connecting them to the underlying tissue microstructure (Wu et al., 2022, Sandgren et al., 2021, Zhang et al., 2020).

## Conclusions

This study introduces restriction-weighted q-space trajectory imaging (ResQ); a dMRI framework that enables a joint investigation of restriction effects and tensor-valued diffusion encoding. Using a longitudinal study of human prostate cancer xenografts, we demonstrated that restrictions have a substantial influence on the signal and derived parameters when using a range of gradient waveforms. ResQ successfully captured these signal dynamics and enabled estimation of both conventional diffusion parameters and their dependence on restriction weighting.

ResQ revealed pronounced microstructure-sensitive diffusion changes associated with tumor development and radiotherapy. Treatment effects were detectable within one day and were primarily reflected in parameters related to diffusivity (*D* and Δ*D*) and isotropic diffusional variance (*V*_Di_), indicating sensitivity of the method to therapy-induced alterations in tissue microstructure. These findings demonstrate that restriction-weighted tensor-valued diffusion encoding can provide complementary information to conventional diffusion MRI approaches.

Our study highlights the critical importance of gradient waveform design for accurate signal representations and biophysical modeling, particularly in tissues that exhibit restricted diffusion. Failing to account for such effects can introduce gross bias in the estimated diffusion parameters and subsequent interpretation, seen especially as an overestimation of microscopic diffusion anisotropy. However, the magnitude and direction of the bias are not universal but rather depend on the experimental configuration and the tissue microstructure, warranting careful evaluation of restriction-weighting effects in any given experimental setting.

We have shared open-source tools for experimental design and analysis to enable continued development of ResQ. Future studies should evaluate ResQ in human prostate imaging and across additional tumor types to establish its potential for clinical translation and for developing microstructure-sensitive biomarkers in oncologic imaging and beyond.

## Data availability statement

MRI data is available upon reasonable request from the corresponding author. The modified waveform optimizer is available in open source at github.com/filip-szczepankiewicz/NOW. Code for ResQ parameter estimation is available at github.com/filip-szczepankiewicz/ResQ. Resources related specifically to this study, such as gradient waveform files, detailed experimental descriptions, and processing code are available at github.com/filip-szczepankiewicz/Szczepankiewicz_NMRBiomed_2026.

## Acknowledgements

We thank Mathew Budde for generously providing the Bruker pulse sequence which can be found at https://github.com/mdbudde/mcw_Preclinical_MRIsequences. We thank Susanne Geres and Lund University Bioimaging Center (LBIC) for furnishing the animal care and experimental resources.

## Author contribution statement

Filip Szczepankiewicz: conceptualization; methodology; software; formal analysis; investigation; resources; data curation; visualization; supervision; writing – original draft; writing – review & editing; project administration; funding acquisition. Malwina Molendowska: data curation; investigation; writing – review & editing. Samo Lasič: methodology; writing – review & editing. Marcella E Safi: methodology; writing – review & editing. Michael Gottschalk: methodology; data curation; writing – review & editing. Evangelia Sereti: writing – review & editing. Anders Bjartell: writing – review & editing. Linda Knutsson: resources; writing – review & editing. Oskar Vilhelmsson Timmermand: methodology; investigation; writing – review & editing. Crister Ceberg: resources; methodology; writing – review & editing. Joanna Strand: conceptualization; methodology; investigation; resources; data curation; writing – review & editing; supervision; project administration; funding acquisition.

## List of abbreviations

DKI: Diffusional kurtosis imaging
dMRI: Diffusion MRI
DTI: Diffusion tensor imaging
DWI: Diffusion-weighted imaging
IQR: Interquartile range
LNCaP: Lymph node carcinoma of the prostate
LTE: Linear b-tensor encoding
MRI: Magnetic resonance imaging
PI-RADS: Prostate Imaging – Reporting and Data System
PSA: Prostate-specific antigen
PSMA: Prostate-specific membrane antigen
PTE: Planar b-tensor encoding
QTI: Q-space trajectory imaging
ResQ: Restriction-weighted q-space trajectory imaging
RT: Radiotherapy
SNR: Signal-to-noise ratio
STE: Spherical b-tensor encoding

## Footnotes

1 Note that the factor 3 is replaced by 9/2 if we were to consider the sample variance rather than the population variance.

## References

Agrotis, G., Pooch, E., Abdelatty, M., Benson, S., Vassiou, A., Vlychou, M., Beets-Tan, R. G. H. & Schoots, I. G. 2025. Diagnostic performance of ADC and ADCratio in MRI-based prostate cancer assessment: A systematic review and meta-analysis. Eur Radiol, 35, 404–416.

Almansour, H., Schick, F., Nachbar, M., Afat, S., Fritz, V., Thorwarth, D., Zips, D., Bertram, F., Muller, A. C., Nikolaou, K., Othman, A. E. & Wegener, D. 2023. Longitudinal monitoring of Apparent Diffusion Coefficient (ADC) in patients with prostate cancer undergoing MR-guided radiotherapy on an MR-Linac at 1.5 T: a prospective feasibility study. Radiol Oncol, 57, 184–190.

Alves, R., Henriques, R. N., Kerkela, L., Chavarrias, C., Jespersen, S. N. & Shemesh, N. 2022. Correlation Tensor MRI deciphers underlying kurtosis sources in stroke. Neuroimage, 247, 118833.

Bak, M. & Nielsen, N. C. 1997. REPULSION, A Novel Approach to Efficient Powder Averaging in Solid-State NMR. J Magn Reson, 125, 132–9.

Baron, C. A. & Beaulieu, C. 2014. Oscillating gradient spin-echo (OGSE) diffusion tensor imaging of the human brain. Magn Reson Med, 72, 726–36.

Basser, P. J., Mattiello, J. & Le Bihan, D. 1994. MR diffusion tensor spectroscopy and imaging. Biophys J, 66, 259–67.

Benjamini, D. & Basser, P. J. 2020. Multidimensional correlation MRI. NMR Biomed, e4226.

Bigun, J. optimal orientation detection of linear symmetry. Proc. of the IEEE First international conference on computer vision, June 8–11 1987 London. 433–438.

Bischoff, L. M., Endler, C., Krausewitz, P., Ellinger, J., Klumper, N., Isaak, A., Mesropyan, N., Kravchenko, D., Nowak, S., Kuetting, D., Sprinkart, A. M., Murtz, P., Pieper, C. C., Attenberger, U. & Luetkens, J. A. 2024. Ultra-high gradient performance 3-Tesla MRI for super-fast and high-quality prostate imaging: initial experience. Insights Imaging, 15, 287.

Bourne, R. M., Bongers, A., Chatterjee, A., Sved, P. & Watson, G. 2016. Diffusion anisotropy in fresh and fixed prostate tissue ex vivo. Magn Reson Med, 76, 626–34.

Bourne, R. M., Kurniawan, N., Cowin, G., Sved, P. & Watson, G. 2012. Microscopic diffusion anisotropy in formalin fixed prostate tissue: preliminary findings. Magn Reson Med, 68, 1943–8.

Brabec, J., Szczepankiewicz, F., Lennartsson, F., Englund, E., Pebdani, H., Bengzon, J., Knutsson, L., Westin, C. F., Sundgren, P. C. & Nilsson, M. 2022. Histogram analysis of tensor-valued diffusion MRI in meningiomas: Relation to consistency, histological grade and type. Neuroimage Clin, 33, 102912.

Chakwizira, A., Szczepankiewicz, F. & Nilsson, M. 2025. Diffusion MRI with double diffusion encoding and variable mixing times disentangles water exchange from transient kurtosis. Sci Rep, 15, 8747.

Chakwizira, A., Westin, C. F., Brabec, J., Lasic, S., Knutsson, L., Szczepankiewicz, F. & Nilsson, M. 2022. Diffusion MRI with pulsed and free gradient waveforms: Effects of restricted diffusion and exchange. NMR Biomed, e4827.

Chatterjee, A., Bourne, R. M., Wang, S., Devaraj, A., Gallan, A. J., Antic, T., Karczmar, G. S. & Oto, A. 2018. Diagnosis of Prostate Cancer with Noninvasive Estimation of Prostate Tissue Composition by Using Hybrid Multidimensional MR Imaging. Radiology, 287, 864–873.

Chatterjee, A. & Dwivedi, D. K. 2024. MRI-based virtual pathology of the prostate. MAGMA, 37, 709–720.

Chatterjee, A., Gallan, A., Fan, X., Medved, M., Akurati, P., Bourne, R. M., Antic, T., Karczmar, G. S. & Oto, A. 2023. Prostate Cancers Invisible on Multiparametric MRI: Pathologic Features in Correlation with Whole-Mount Prostatectomy. Cancers (Basel), 15.

Cho, E., Baek, H. J., Szczepankiewicz, F., An, H. J. & Jung, E. J. 2024. Imaging evaluation focused on microstructural tissue changes using tensor-valued diffusion encoding in breast cancers after neoadjuvant chemotherapy: is it a promising way forward? Gland Surg, 13, 1387–1399.

Cifuentes, E., Croxen, R., Menon, M., Barrack, E. R. & Reddy, G. P. 2003. Synchronized prostate cancer cells for studying androgen regulated events in cell cycle progression from G1 into S phase. J Cell Physiol, 195, 337–45.

Cordero-Grande, L., Christiaens, D., Hutter, J., Price, A. N. & Hajnal, J. V. 2019. Complex diffusion-weighted image estimation via matrix recovery under general noise models. Neuroimage, 200, 391–404.

Cory, D. G., Garroway, A. N. & Miller, J. B. 1990. Applications of Spin Transport as a Probe of Local Geometry. Abstracts of Papers of the American Chemical Society, 199, 105.

De Almeida Martins, J. P. & Topgaard, D. 2018. Multidimensional correlation of nuclear relaxation rates and diffusion tensors for model-free investigations of heterogeneous anisotropic porous materials. Sci Rep, 8, 2488.

De Swiet, T. M. & Mitra, P. P. 1996. Possible Systematic Errors in Single-Shot Measurements of the Trace of the Diffusion Tensor. J Magn Reson B, 111, 15–22.

Does, M. D., Parsons, E. C. & Gore, J. C. 2003. Oscillating gradient measurements of water diffusion in normal and globally ischemic rat brain. Magn Reson Med, 49, 206–15.

Epstein, C. L. & Schotland, J. 2008. The bad truth about Laplace’s transform. Siam Review, 50, 504–520.

Eriksson, S., Lasič, S., Nilsson, M., Westin, C. F. & Topgaard, D. 2015. NMR diffusion-encoding with axial symmetry and variable anisotropy: Distinguishing between prolate and oblate microscopic diffusion tensors with unknown orientation distribution. J Chem Phys, 142, 104201.

Finsterbusch, J. 2011. Multiple-Wave-Vector Diffusion-Weighted NMR. Annual Reports on NMR Spectroscopy, 72, 225–299.

Fokkinga, E., Hernandez-Tamames, J. A., Ianus, A., Nilsson, M., Tax, C. M. W., Perez-Lopez, R. & Grussu, F. 2023. Advanced Diffusion-Weighted MRI for Cancer Microstructure Assessment in Body Imaging, and Its Relationship With Histology. J Magn Reson Imaging.

Foltz, W. D., Wu, A., Chung, P., Catton, C., Bayley, A., Milosevic, M., Bristow, R., Warde, P., Simeonov, A., Jaffray, D. A., Haider, M. A. & Menard, C. 2013. Changes in apparent diffusion coefficient and T2 relaxation during radiotherapy for prostate cancer. J Magn Reson Imaging, 37, 909–16.

Grebenkov, D. S. 2007. NMR survey of reflected Brownian motion. Reviews of Modern Physics, 79, 1077–1137.

Gudbjartsson, H. & Patz, S. 1995. The Rician Distribution of Noisy MRI Data. Magn Reson Imaging, 34, 910–914.

Hectors, S. J., Semaan, S., Song, C., Lewis, S., Haines, G. K., Tewari, A., Rastinehad, A. R. & Taouli, B. 2018. Advanced Diffusion-weighted Imaging Modeling for Prostate Cancer Characterization: Correlation with Quantitative Histopathologic Tumor Tissue Composition-A Hypothesis-generating Study. Radiology, 286, 918–928.

Henriques, R. N., Jespersen, S. N. & Shemesh, N. 2020. Correlation tensor magnetic resonance imaging. Neuroimage, 211, 116605.

Henriques, R. N., Jespersen, S. N. & Shemesh, N. 2021. Evidence for microscopic kurtosis in neural tissue revealed by correlation tensor MRI. Magn Reson Med, 86, 3111–3130.

Jensen, J. H., Helpern, J. A., Ramani, A., Lu, H. & Kaczynski, K. 2005. Diffusional kurtosis imaging: the quantification of non-gaussian water diffusion by means of magnetic resonance imaging. Magn Reson Med, 53, 1432–40.

Jespersen, S. N. 2025. Isotropic sampling of tensor-encoded diffusion MRI. Magn Reson Med, 93, 2040–2048.

Jespersen, S. N., Lundell, H., Sønderby, C. K. & Dyrby, T. B. 2013. Orientationally invariant metrics of apparent compartment eccentricity from double pulsed field gradient diffusion experiments. NMR Biomed, 26, 1647–62.

Jespersen, S. N., Olesen, J. L., Ianus, A. & Shemesh, N. 2017. Anisotropy in “isotropic diffusion” measurements due to nongaussian diffusion. 1712.02290v1 [physics.bio-ph].

Jespersen, S. N., Olesen, J. L., Ianus, A. & Shemesh, N. Implications of nongaussian diffusion on the interpretation of multidimensional diffusion measurements. Proc. Intl. Soc. Mag. Reson. Med. 26, 2018. 0886.

Jiang, H., Svenningsson, L. & Topgaard, D. 2023. Multidimensional encoding of restricted and anisotropic diffusion by double rotation of the q vector. Magn Reson (Gott), 4, 73–85.

Jiang, X., Li, H., Xie, J., Mckinley, E. T., Zhao, P., Gore, J. C. & Xu, J. 2017. In vivo imaging of cancer cell size and cellularity using temporal diffusion spectroscopy. Magn Reson Med, 78, 156–164.

Johnson, J. T. E., Irfanoglu, M. O., Manninen, E., Ross, T. J., Yang, Y., Laun, F. B., Martin, J., Topgaard, D. & Benjamini, D. 2024. In vivo disentanglement of diffusion frequency-dependence, tensor shape, and relaxation using multidimensional MRI. Hum Brain Mapp, 45, e26697.

Johnston, E. W., Bonet-Carne, E., Ferizi, U., Yvernault, B., Pye, H., Patel, D., Clemente, J., Piga, W., Heavey, S., Sidhu, H. S., Giganti, F., O’Callaghan, J., Brizmohun Appayya, M., Grey, A., Saborowska, A., Ourselin, S., Hawkes, D., Moore, C. M., Emberton, M., Ahmed, H. U., Whitaker, H., Rodriguez-Justo, M., Freeman, A., Atkinson, D., Alexander, D., Panagiotaki, E. & Punwani, S. 2019. VERDICT MRI for Prostate Cancer: Intracellular Volume Fraction versus Apparent Diffusion Coefficient. Radiology, 291, 391–397.

Jones, D. K., Horsfield, M. A. & Simmons, A. 1999. Optimal strategies for measuring diffusion in anisotropic systems by magnetic resonance imaging. Magn Reson Med, 42, 515–25.

Kiselev, V. 2011. The cumulant expansion: an overarching mathematical framework for understanding diffusion NMR. In: Jones, D. K. (ed.) Diffusion MRI: Theory, Methods, and Applications.

Klein, S., Staring, M., Murphy, K., Viergever, M. A. & Pluim, J. P. 2010. elastix: a toolbox for intensity-based medical image registration. IEEE Trans Med Imaging, 29, 196–205.

Lampinen, B., Szczepankiewicz, F., Latt, J., Knutsson, L., Martensson, J., Bjorkman-Burtscher, I. M., Van Westen, D., Sundgren, P. C., Stahlberg, F. & Nilsson, M. 2023. Probing brain tissue microstructure with MRI: principles, challenges, and the role of multidimensional diffusion-relaxation encoding. Neuroimage, 282, 120338.

Langbein, B. J., Szczepankiewicz, F., Westin, C. F., Bay, C., Maier, S. E., Kibel, A. S., Tempany, C. M. & Fennessy, F. M. 2021. A Pilot Study of Multidimensional Diffusion MRI for Assessment of Tissue Heterogeneity in Prostate Cancer. Invest Radiol, 56, 845–853.

Lasič, S., Just, N., Nilsson, M., Szczepankiewicz, F., Budde, M. & Lundell, H. 2025. Spectral principal axis system (SPAS) and tuning of tensor-valued encoding for microscopic anisotropy and time-dependent diffusion in the rat brain. Imaging Neuroscience, 3.

Lasič, S., Szczepankiewicz, F., Eriksson, S., Nilsson, M. & Topgaard, D. 2014. Microanisotropy imaging: quantification of microscopic diffusion anisotropy and orientational order parameter by diffusion MRI with magic-angle spinning of the q-vector. Frontiers in Physics, 2, 11.

Lemberskiy, G., Rosenkrantz, A. B., Veraart, J., Taneja, S. S., Novikov, D. S. & Fieremans, E. 2017. Time-Dependent Diffusion in Prostate Cancer. Invest Radiol, 52, 405–411.

Lundell, H. & Lasič, S. 2020. Diffusion Encoding with General Gradient Waveforms. In: Topgaard, D. (ed.) Advanced Diffusion Encoding Methods in MRI. Royal Society of Chemistry, 2020.

Lundell, H., Nilsson, M., Dyrby, T. B., Parker, G. J. M., Cristinacce, P. L. H., Zhou, F. L., Topgaard, D. & Lasič, S. 2019. Multidimensional diffusion MRI with spectrally modulated gradients reveals unprecedented microstructural detail. Sci Rep, 9, 9026.

Ma, X., Seres, P., Kinnaird, A., Fung, C., Feiweier, T. & Beaulieu, C. 2025. Diffusion time effects over the adult lifespan indicates persistent zone-specific microstructural alterations in the human prostate with aging. Magn Reson Med, 93, 2059–2069.

Molendowska, M., Engel, M., Mueller, L., Lasič, S., Jones, D. K., Tax, C. M. & Szczepankiewicz, F. High Fidelity Imaging of Tissue Heterogeneity, Micro-Anisotropy and Diffusion-Time Effects in Prostate Cancer. Proc. Int. Soc. Magn. Reson. Med. 32, 2024 Singapore.

Molendowska, M., Fasano, F., Rudrapatna, U., Kimmlingen, R., Jones, D. K., Kusmia, S., Tax, C. M. W. & Evans, C. J. 2022. Physiological effects of human body imaging with 300 mT/m gradients. Magn Reson Med, 87, 2512–2520.

Molendowska, M., Lasič, S., Strand, J. & Szczepankiewicz, F. A novel framework for restriction-weighted q-space trajectory imaging (resQ) demonstrated in prostate cancer. ISMRM workshop on 40 Years of Diffusion: Past, Present & Future Perspectives, 2025a Kyoto, Japan.

Molendowska, M., Olsson, V., Mortensen, F., Testud, F. & Szczepankiewicz, F. A stimulation-adapted gradient design for velocity-compensated IVIM demonstrated in prostate at 200 mT/m. 40 Years of Diffusion: Past, Present & Future Perspectives, 2025b Kyoto, Japan. Int. Soc. Magn. Reson. Med.

Möller, T. & Hughes, J. F. 1999. Efficiently building a matrix to rotate one vector to another.

Nam, R., Patel, C., Milot, L., Hird, A., Wallis, C., Macinnis, P., Singh, M., Emmenegger, U., Sherman, C. & Haider, M. A. 2022. Prostate MRI versus PSA screening for prostate cancer detection (the MVP Study): a randomised clinical trial. BMJ Open, 12, e059482.

Narvaez, O., Svenningsson, L., Yon, M., Sierra, A. & Topgaard, D. 2022. Massively Multidimensional Diffusion-Relaxation Correlation MRI. Frontiers in Physics, 9.

Nilsson, M., Eklund, G., Szczepankiewicz, F., Skorpil, M., Bryskhe, K., Westin, C. F., Lindh, C., Blomqvist, L. & Jaderling, F. 2021. Mapping prostatic microscopic anisotropy using linear and spherical b-tensor encoding: A preliminary study. Magn Reson Med, 86, 2025–2033.

Nilsson, M., Lasic, S., Drobnjak, I., Topgaard, D. & Westin, C. F. 2017. Resolution limit of cylinder diameter estimation by diffusion MRI: The impact of gradient waveform and orientation dispersion. NMR Biomed, 30.

Nilsson, M., Szczepankiewicz, F., Van Westen, D. & Hansson, O. 2015. Extrapolation-Based References Improve Motion and Eddy-Current Correction of High B-Value DWI Data: Application in Parkinson’s Disease Dementia. PLoS One, 10, e0141825.

Novello, L., Henriques, R. N., Ianus, A., Feiweier, T., Shemesh, N. & Jovicich, J. 2022. In vivo Correlation Tensor MRI reveals microscopic kurtosis in the human brain on a clinical 3T scanner. Neuroimage, 254, 119137.

Novikov, D. S., Fieremans, E., Jensen, J. H. & Helpern, J. A. 2011. Random walk with barriers. Nat Phys, 7, 508–514.

Novikov, D. S., Jensen, J. H., Helpern, J. A. & Fieremans, E. 2014. Revealing mesoscopic structural universality with diffusion. Proc Natl Acad Sci U S A, 111, 5088–93.

Novikov, D. S., Kiselev, V. G. & Jespersen, S. N. 2018. On modeling. Magn Reson Med, 79, 3172–3193.

Padhani, A. R., Haider, M. A., Villers, A. & Barentsz, J. O. 2019. Multiparametric Magnetic Resonance Imaging for Prostate Cancer Detection: What We See and What We Miss. Eur Urol, 75, 721–722.

Panagiotaki, E. & Alexander, D. C. 2014. Non-invasive quantification of solid tumor microstructure using VERDICT MRI.

Pasquier, D., Hadj Henni, A., Escande, A., Tresch, E., Reynaert, N., Colot, O., Lartigau, E. & Betrouni, N. 2018. Diffusion weighted MRI as an early predictor of tumor response to hypofractionated stereotactic boost for prostate cancer. Sci Rep, 8, 10407.

Ronen, I., Moeller, S., Ugurbil, K. & Kim, D. S. 2006. Analysis of the distribution of diffusion coefficients in cat brain at 9.4 T using the inverse Laplace transformation. Magn Reson Imaging, 24, 61–8.

Roth, D., Safi, M., Vilhelmsson Timmermand, O., Sereti, E., Molendowska, M., Gottschalk, M., Bjartell, A., Ceberg, C., Szczepankiewicz, F. & Strand, J. 2024. Evaluation of superficial xenograft volume estimation by ultrasound and caliper against MRI in a longitudinal pre-clinical radiotherapeutic setting. PLoS One, 19, e0307558.

Sandgren, K., Nilsson, E., Keeratijarut Lindberg, A., Strandberg, S., Blomqvist, L., Bergh, A., Friedrich, B., Axelsson, J., Ogren, M., Ogren, M., Widmark, A., Thellenberg Karlsson, C., Soderkvist, K., Riklund, K., Jonsson, J. & Nyholm, T. 2021. Registration of histopathology to magnetic resonance imaging of prostate cancer. Phys Imaging Radiat Oncol, 18, 19–25.

Shemesh, N., Ozarslan, E., Adiri, T., Basser, P. J. & Cohen, Y. 2010. Noninvasive bipolar double-pulsed-field-gradient NMR reveals signatures for pore size and shape in polydisperse, randomly oriented, inhomogeneous porous media. J Chem Phys, 133, 044705.

Singh, S., Rogers, H., Kanber, B., Clemente, J., Pye, H., Johnston, E. W., Parry, T., Grey, A., Dinneen, E., Shaw, G., Heavey, S., Stopka-Farooqui, U., Haider, A., Freeman, A., Giganti, F., Atkinson, D., Moore, C. M., Whitaker, H. C., Alexander, D. C., Panagiotaki, E. & Punwani, S. 2022. Avoiding Unnecessary Biopsy after Multiparametric Prostate MRI with VERDICT Analysis: The INNOVATE Study. Radiology, 212536.

Sjölund, J., Szczepankiewicz, F., Nilsson, M., Topgaard, D., Westin, C. F. & Knutsson, H. 2015. Constrained optimization of gradient waveforms for generalized diffusion encoding. J Magn Reson, 261, 157–168.

Song, I., Kim, C. K., Park, B. K. & Park, W. 2010. Assessment of response to radiotherapy for prostate cancer: value of diffusion-weighted MRI at 3 T. AJR Am J Roentgenol, 194, W477– 82.

Stejskal, E. O. & Tanner, J. E. 1965. Spin Diffusion Measurement: Spin echoes in the Presence of a Time-Dependent Field Gradient. the journal of chemical physics, 42, 288–292.

Stepisnik, J. 1993. Time-Dependent Self-Diffusion by Nmr Spin-Echo. Physica B, 183, 343–350.

Stepisnik, J. 1999. Validity limits of Gaussian approximation in cumulant expansion for diffusion attenuation of spin echo. Physica B, 270, 110–117.

Stoyanova, R., Chinea, F., Kwon, D., Reis, I. M., Tschudi, Y., Parra, N. A., Breto, A. L., Padgett, K. R., Dal Pra, A., Abramowitz, M. C., Kryvenko, O. N., Punnen, S. & Pollack, A. 2018. An Automated Multiparametric MRI Quantitative Imaging Prostate Habitat Risk Scoring System for Defining External Beam Radiation Therapy Boost Volumes. Int J Radiat Oncol Biol Phys, 102, 821–829.

Szczepankiewicz, F., Safi, M., Ceberg, C., Gottschalk, M., Sereti, E., Bjartell, A., Vilhelmsson Timmermand, O., Knutsson, L.Strand, S.-E. & Strand, J. Multidimensional diffusion MRI for monitoring radiotherapy response in human prostate cancer xenografts in mice: a longitudinal pilot study (4157). Proc Int Soc Magn Reson Med, 2023 Toronto, ON, Canada.

Szczepankiewicz, F., Sjölund, J., Ståhlberg, F., Lätt, J. & Nilsson, M. 2019a. Tensor-valued diffusion encoding for diffusional variance decomposition (DIVIDE): Technical feasibility in clinical MRI systems. PLoS One, 14, e0214238.

Szczepankiewicz, F., Van Westen, D., Englund, E., Westin, C. F., Stahlberg, F., Latt, J., Sundgren, P. C. & Nilsson, M. 2016. The link between diffusion MRI and tumor heterogeneity: Mapping cell eccentricity and density by diffusional variance decomposition (DIVIDE). Neuroimage, 142, 522–532.

Szczepankiewicz, F., Westin, C. F. & Nilsson, M. 2019b. Maxwell-compensated design of asymmetric gradient waveforms for tensor-valued diffusion encoding. Magn Reson Med.

Szczepankiewicz, F., Westin, C. F. & Nilsson, M. 2021. Gradient waveform design for tensor-valued encoding in diffusion MRI. J Neurosci Methods, 109007.

Tempany, C. M., Jayender, J., Kapur, T., Bueno, R., Golby, A., Agar, N. & Jolesz, F. A. 2015. Multimodal imaging for improved diagnosis and treatment of cancers. Cancer, 121, 817–27.

Topgaard, D. 2016. Chapter 7. NMR Methods for Studying Microscopic Diffusion Anisotropy. In: Valiullin, R. (ed.) Diffusion NMR of Confined Systems. Royal Society of Chemistry, Cambridge, UK.

Topgaard, D. 2025. Validity of the Gaussian phase distribution approximation for analysis of isotropic diffusion encoding applied to restricted diffusion in a cylinder. Magn Reson Lett, 5, 200196.

Tournier, J. D., Calamante, F. & Connelly, A. 2012. MRtrix: Diffusion tractography in crossing fiber regions. International Journal of Imaging Systems and Technology, 22, 53–66.

Wang, S., Peng, Y., Medved, M., Yousuf, A. N., Ivancevic, M. K., Karademir, I., Jiang, Y., Antic, T., Sammet, S., Oto, A. & Karczmar, G. S. 2014. Hybrid multidimensional T(2) and diffusion-weighted MRI for prostate cancer detection. J Magn Reson Imaging, 39, 781–8.

Weinreb, J. C., Barentsz, J. O., Choyke, P. L., Cornud, F., Haider, M. A., Macura, K. J., Margolis, D., Schnall, M. D., Shtern, F., Tempany, C. M., Thoeny, H. C. & Verma, S. 2016. PI-RADS Prostate Imaging - Reporting and Data System: 2015, Version 2. Eur Urol, 69, 16–40.

Westin, C. F., Knutsson, H., Pasternak, O., Szczepankiewicz, F., Özarslan, E., Van Westen, D., Mattisson, C., Bogren, M., O’Donnell, L. J., Kubicki, M., Topgaard, D. & Nilsson, M. 2016. Q-space trajectory imaging for multidimensional diffusion MRI of the human brain. Neuroimage, 135, 345–62.

Wu, D., Jiang, K., Li, H., Zhang, Z., Ba, R., Zhang, Y., Hsu, Y. C., Sun, Y. & Zhang, Y. D. 2022. Time-Dependent Diffusion MRI for Quantitative Microstructural Mapping of Prostate Cancer. Radiology, 303, 578–587.

Yadav, S. S., Stockert, J. A., Hackert, V., Yadav, K. K. & Tewari, A. K. 2018. Intratumor heterogeneity in prostate cancer. Urol Oncol, 36, 349–360.

Yeh, F. C. 2022. Population-based tract-to-region connectome of the human brain and its hierarchical topology. Nat Commun, 13, 4933.

Zhang, Z., Wu, H. H., Priester, A., Magyar, C., Afshari Mirak, S., Shakeri, S., Mohammadian Bajgiran, A., Hosseiny, M., Azadikhah, A., Sung, K., Reiter, R. E., Sisk, A. E., Raman, S. & Enzmann, D. R. 2020. Prostate Microstructure in Prostate Cancer Using 3-T MRI with Diffusion-Relaxation Correlation Spectrum Imaging: Validation with Whole-Mount Digital Histopathology. Radiology, 296, 348–355.

Zhu, A., Tarasek, M., Hua, Y., Fiveland, E., Maier, S. E., Mazaheri, Y., Fung, M., Westin, C. F., Yeo, D. T. B., Szczepankiewicz, F., Tempany, C., Akin, O. & Foo, T. K. F. 2024. Human prostate MRI at ultrahigh-performance gradient: A feasibility study. Magn Reson Med, 91, 640–648.

