## Supplement for "Restriction-weighted q-space trajectory imaging (ResQ): Toward mapping diffusion-time effects with tensor-valued diffusion encoding in human prostate cancer xenografts"

### Supplementary materials

To gauge data quality, we provide examples of data from a representative examination (Figure S1), i.e., the one that had an average SNR (within the examination) that was closest to the global average (across all examinations). Additionally, we provide an overview of the effect of denoising by showing the difference between single images before and after denoising.

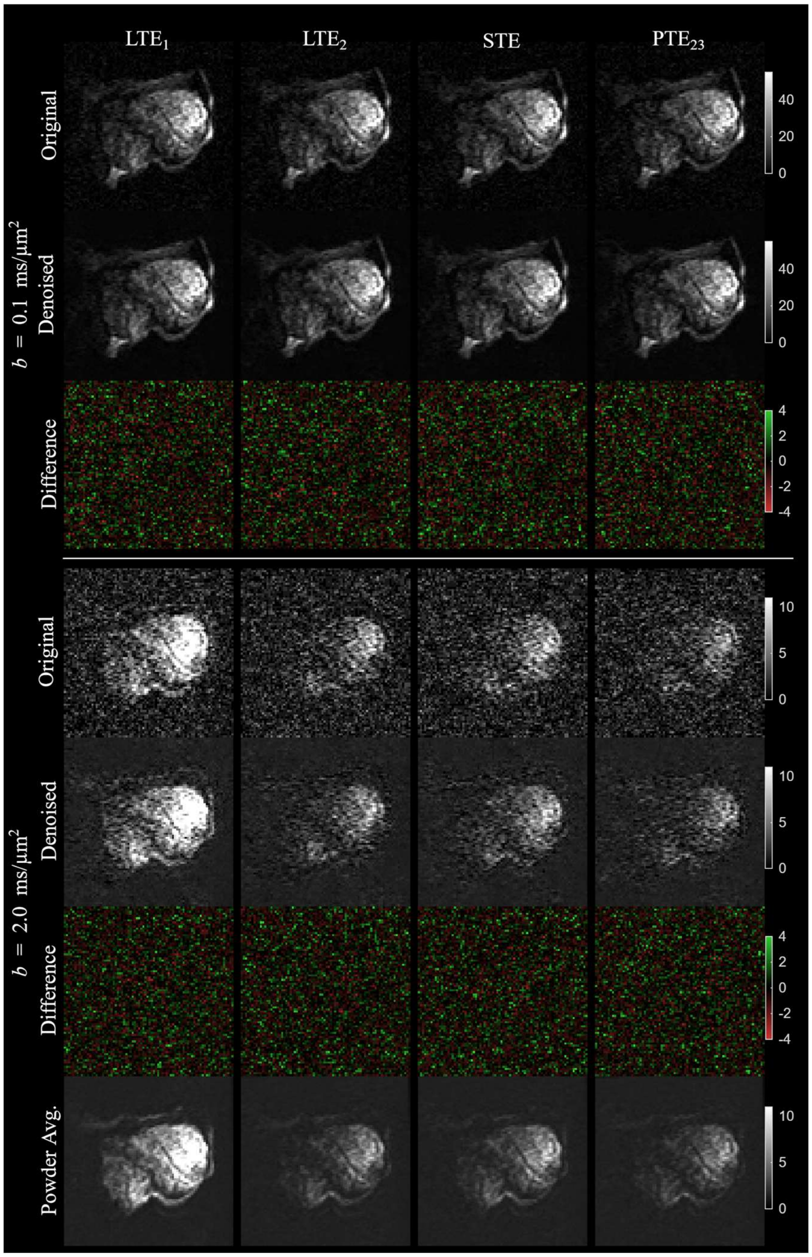


Figure S1 – Top and bottom sections show single acquisitions at low and high b-values along with the denoised image and difference between them. The lack of structure in the difference maps indicates that the denoising did not remove true signal components that could otherwise affect the accuracy of parameter estimation. The bottom row shows the powder-averaged signal at the highest b-value after denoising. This shows the quality of the data at the lowest SNR within the given data set. It is also clearly visible that the signal from $LTE_1$—the waveform with the lowest frequencies and longest diffusion time—has a markedly higher signal than the other waveform variants.

We also provide a more complete overview of the structural anisotropy maps calculated by structure tensor analysis in Figure S2. Generally, structural anisotropy was low, and the samples did not exhibit convincingly anisotropic structures even in the regions where SA was highest.

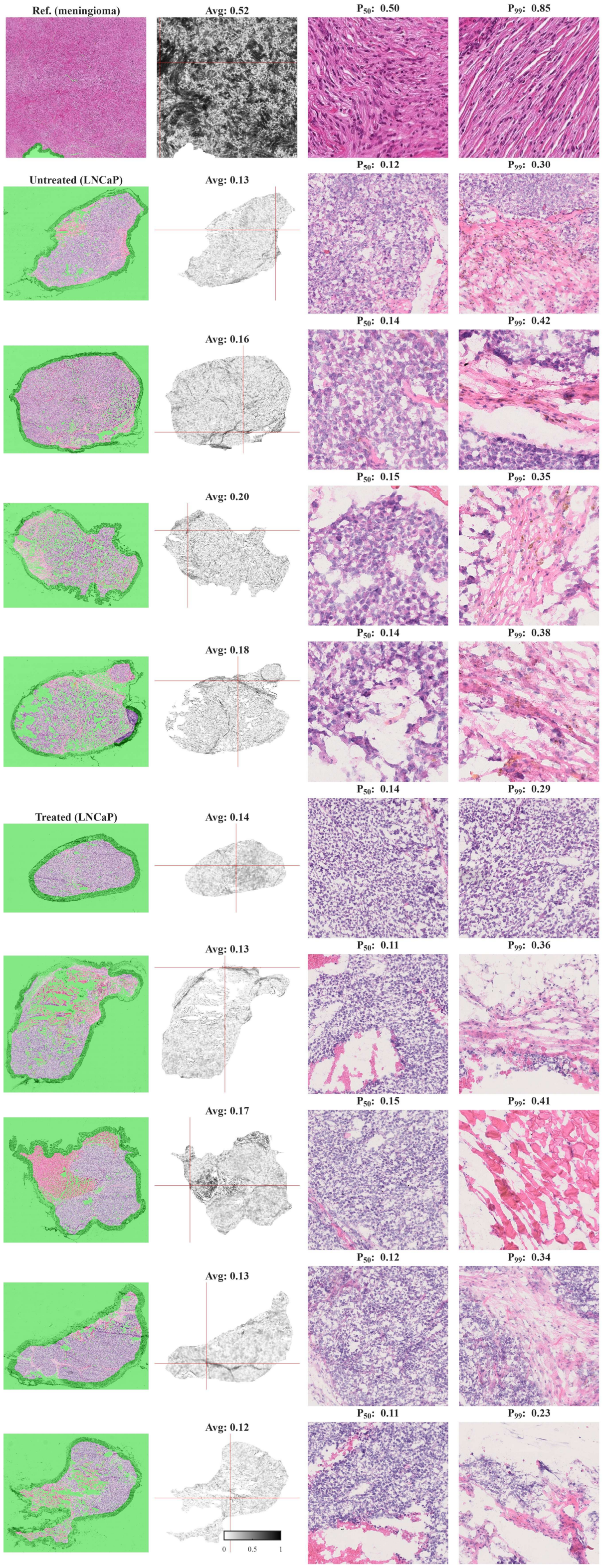


Figure S2 – Histology with H&E staining (1st column) and structural anisotropy (SA) maps from structure tensor analysis (2nd column). Top row shows a fibroblastic meningioma with high anisotropy for reference (Szczepankiewicz et al. 2016); remaining rows are LNCaP tumors from this study. Rightmost columns show regions that have median SA (50th percentile, $P_{50}$) and close to the highest SA (99th percentile, $P_{99}$). The green area shows the masking used to remove large empty spaces and edges which otherwise biased the analysis. Red crosshair in SA maps shows the position of the zoomed in region that corresponds to $P_{99}$. The highest SA appears in tissue transitions or rare patches of stroma. The width of the magnified patches is approximately 300 µm.
